# Soil nitrogen dynamics in riparian zones of two Arctic rivers

**DOI:** 10.64898/2026.10.12.758441

**Authors:** Wasi Hashmi, Marit Ebbinghaus, Meret Carstensen, Anna Wolgast, Carlos Palacin-Lizarbe, Jenie Gil, Lukas Kohl, Jukka Pumpanen, Tina Sanders, Henri M.P. Siljanen, Claudia Fiencke, Maija E. Marushchak

## Abstract

Riparian zones act as protective buffers, filtering land-to-river nutrient inputs through retention and nitrogen (N) gas losses. While rapid Arctic warming and permafrost thaw may enhance terrestrial nutrient inputs to aquatic systems, comprehensive assessments of N cycling in Arctic riparian soils are lacking. Conventionally, these zones are considered inactive due to N limitation of Arctic soils and low temperatures, leaving their N cycling unexplored. To address this, we investigated N pools and turnover processes in riparian zones of two Arctic rivers in Finland and Greenland. We combined measurements of the extractable N pool and *in-situ* nitrous oxide (N_2_O) flux with ^15^N-labeling techniques to quantify gross mineralization, nitrification, potential denitrification, and potential dissimilatory nitrate reduction to ammonium (DNRA).

Our findings reveal that mineral N cycling in high-latitude riparian zones is more active than previously assumed. Despite low temperatures and low to intermediate soil organic carbon content, we found high gross N mineralization rates, alongside detectable rates of gross nitrification and potential DNRA. We attribute the unexpectedly high N activity in C-poor soils to low C/N ratios, which create a stoichiometric imbalance, thereby enhancing N mineralization. The presence or absence of vegetation largely determined the dominance and magnitude of these pathways. While dense vascular plant cover was associated with higher extractable N pools and gross mineralization rate, low nitrification rates and elevated DNRA potential indicated a tight N cycle. Conversely, sparsely vegetated patches with cryptogamic biocrusts acted as N_2_O hotspots. In bare sandy soils adjacent to the river, we found high gross nitrification rate and detectable N_2_O losses, possibly driven by riverine organic N inputs. Taken together, this research shows that Arctic riparian zones may play an important role in reducing terrestrial N inputs to the rivers, which should be considered when studying the fate of N in the rapidly changing Arctic.

## 1. Introduction

Rapid warming is transforming the Arctic at a rate nearly four times the global average (Rantanen et al., 2022). This thermal amplification is driving widespread permafrost thaw, resulting in deeper active layers and fundamental alteration of hydrological connectivity in high-latitude landscapes (Walvoord & Kurylyk, 2016). As frozen soils thaw, vast stores of sequestered soil organic matter (SOM) and nutrients are being mobilized, increasing the lateral flux of carbon (C) and nitrogen (N) from terrestrial ecosystems into aquatic networks (Turetsky et al., 2020; Tank et al., 2023; Jin et al., 2022). In this context, riparian zones represent dynamic interfaces between land and water which regulate the magnitude, timing and stoichiometry of nutrient delivery to river systems, affecting nutrient transport further downstream to estuaries and, finally, to the ocean (Pinay et al., 2015; Appling et al., 2014). However, in high-latitude riparian zones this filtering capacity is thought to be limited by low temperatures and short growing seasons (Pinay et al. 2018), but very little research is available on N cycling in Arctic riparian soils. Given the observed changes in N transport by large Arctic rivers but limited understanding about how they relate to terrestrial N dynamics (Tank et al., 2023; Sanders et al., 2022; Fuchs et al., 2020; Ruyle et al., 2025), this represents an important research gap.

Within riparian zones, vegetation, SOM content and moisture vary greatly (Fiencke et al., 2022). This may give rise to formation of biogeochemical hotspots, particularly when combined with converging hydrological flow paths that connect substrates from different sources (distinct soil layers, lateral flow, ground water, river water) (Pastor et al., 2020; McClain et al., 2003). In the vicinity of the river soils are commonly bare due to flowing water and fluctuating water level, which causes erosion and hampers the establishment of vegetation cover (Sanders et al., 2022). Absence of plant N uptake enhances the inorganic N availability, increasing the potential for N losses by leaching (Sanders et al., 2010) or as nitrogenous gasses to the atmosphere (Voigt et al., 2020; Walker et al., 2009; Marushchak et al., 2011). However, this is not the only mechanism by which vegetation regulates N availability in the soil. Importantly, plants contribute to SOM accumulation and stabilization by litter inputs and reduction of erosion intensity, thereby enhancing the organic N pool that can be mineralized (Wild et al., 2014). Plants also feed the microbial community with labile C via root exudation, promoting heterotrophic processes, such as denitrification (Henneron et al., 2020; Wild et al., 2014). A critical, yet, often overlooked component of high-latitude vegetation is the biological soil crust (biocrust) composed of lichens, mosses, and cyanobacteria. Biocrusts are the primary ecosystem engineers, which contribute to soil formation and establishment of microbial assembly in organic poor soils, e.g. in glacier forelands (Agnelli et al., 2021; Elbert et al., 2012; Belnap & Lange., 2003), but their role in Arctic riparian soils remains unknown.

The final fate of N in riparian soils — whether it is sequestered, leached, or emitted as gas — depends on the delicate balance between several microbial pathways mediating mineral N transformation. Gross mineralization, the conversion of organic N to ammonium (NH_4_^+^), is the rate-limiting step that largely dictates N availability for organisms (Schimel & Bennett, 2004). Once mineralized, NH_4_^+^ can be immobilized by microbes, taken up by plants, or oxidized to nitrate (NO_3_^−^) in nitrification. Nitrification has great relevance for N losses from pristine northern soils, because it produces easily leachable NO_3_^−^ and nitrous oxide (N_2_O) as a byproduct, and feeds NO_3_^−^ for N_2_O production by denitrification (Voigt et al., 2020, Marushchak et al., 2011). Under anaerobic conditions, commonly occurring in riparian soils, NO_3_^−^ is consumed by two competing pathways: denitrification and dissimilatory nitrate reduction to ammonium (DNRA) (Tiedje et al., 1983; Wang et al., 2020). Denitrification is considered as the dominant pathway in the riparian soils, where it removes bioavailable N from the system by converting NO_3_^−^ to gaseous N_2_O and N_2_ (Palacin-Lizarbe et al., 2019; Pinay et al., 2018). The less studied DNRA process, in contrast, retains N within the ecosystem by converting NO_3_^−^ back into NH_4_^+^, which is less mobile and easily assimilated by plants and microbes (Silver et al., 2001). According to current understanding, the partitioning between these two pathways is controlled by the ratio of available electron donors (C) to electron acceptors (NO_3_^−^) (Pandey et al. 2020). High C/N ratios and strongly reducing conditions generally favour DNRA, while low C/N ratios favour denitrification (Tiedje, 1988; Rütting et al., 2011; Kraft et al 2014). Recent genomic evidence suggests that arctic soils possess higher genetic potential for DNRA relative to denitrification, making arctic tundra exceptional among global biomes (Saghaï et al., 2023), but direct measurements in Arctic riparian soils are lacking.

Despite the importance of mineral N cycling for processing and redistribution of N, empirical data from Arctic riparian zones is scarce (Pastor et al., 2020; Pinay et al., 2018). Most existing studies rely on measurements of net mineralization and/or nitrification rates, which are easy to measure but often not informative (Lupon et al., 2026). Even with low net rates, gross production of NH_4_^+^ or NO_3_^−^ may be substantial if it is matched by equally rapid consumption by immobilization and further mineral N transformation processes. This masks the true intensity of N turnover, as commonly observed in permafrost-affected soils (Ramm et al., 2022), highlighting the importance of measuring gross N rates. However, to the best of our knowledge, there are no thorough investigations available of gross N turnover in Arctic riparian soils, leaving a major gap in the high-latitude biogeochemistry. To address this topic, we investigated soil N dynamics in the riparian zones of two pristine Arctic rivers in Greenland and Finland. We utilized ^15^N stable isotope pool dilution and tracing techniques to quantify gross N transformation rates, including gross mineralization, nitrification, and the rates of potential denitrification and DNRA. Furthermore, spatial variability in soil N pools, *in-situ* GHG fluxes including N_2_O and GHG production in soil incubations was determined. Our aims were to: **(i)** quantify the dominant N pools and turnover processes in Arctic riparian zones, **(ii)** determine their spatial variability in association with soil properties and the presence or absence of vegetation, and **(iii)** determine whether N retention (closed) or loss (leaky) processes dominate in Arctic riparian zones. We hypothesised that Arctic riparian soils are hotspots of N losses and that these hotspots are commonly found in bare soils, where the absence of plant competition for mineral N promotes its loss.

## 2. Methods

### 2.1. Study sites

We studied riparian zones of two pristine Arctic rivers: the Teno River (Finnish: Tenojoki, Norwegian: Tana, Sámi: Deatnu; N 69° 33’ 22’’, E 25° 53’ 35’’, altitude 123 m.a.s.l.) in northern Lapland, Finland and the Red River (Danish: Røde Elv, Greenlandic: Kuussuaq N 69° 16’ 41’’, W 53° 28’ 54’’, altitude 87 m.a.s.l.) in the Kuup Ilua Valley (Danish: Blaesedalen) of Qeqertarsuaq (Disko Island), Greenland (Figure 1a and c). Although both sites are located in the same latitude, the climate is different: The mean annual temperature at the Teno River is −1.1°C and mean annual precipitation 484 mm (years 1991–2020, data from the Utsjoki-Nuorgam weather station, N 70° 04’, E 27° 53’), while the Red River site has colder and less humid climate with mean annual temperature of −3.0°C and the average precipitation 418 mm (years 1991–2017, data from the nearby Arctic station) (Xu & Ambus, 2025).

**Figure 1.**
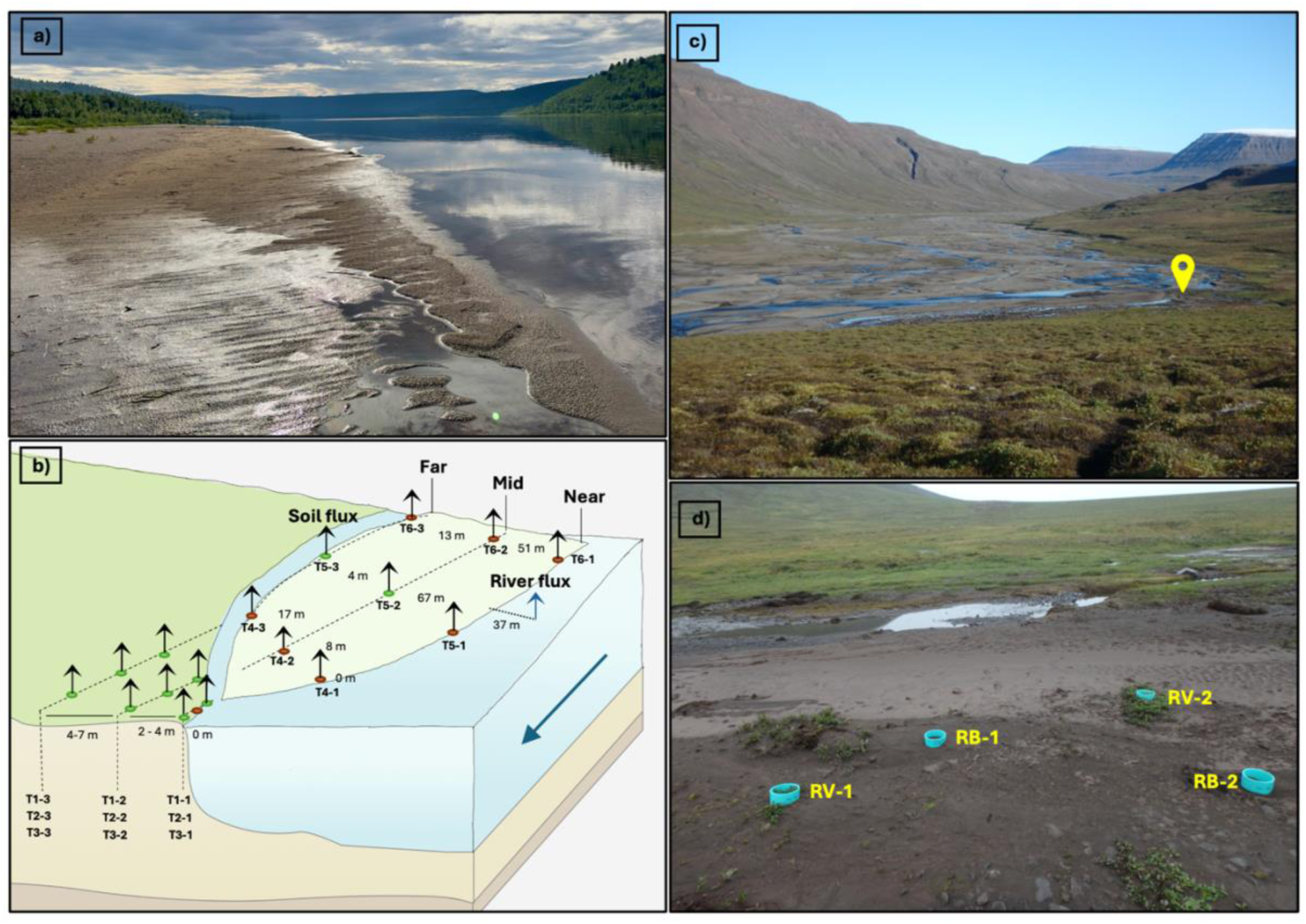
Overview of the study sites. Teno River (Finnish Lapland) during the sampling campaign in August 2023 with a sandbank in the foreground **(a)**, and a schematic diagram of th sampling design **(b)**. Colored dots with arrows show the location of soil sampling and flux measurements, with green and brown dots indicating vegetated and bare plots, respectively. The figure is illustrative and does not represent the actual scale, but approximate distances between the transects (near, mid and far) are given. The blue arrow shows the direction of the river flow. Red River (Disko Island, Greenland) during sampling campaign in September 2023 with th yellow symbol showing the sampling location **(c)**, and sampling plots at the Red River **(d)**.

The Teno River catchment is located in the sporadic permafrost zone and has an area of 16 377 km^2^. It flows for 361 kms at the border between Finland and Norway and discharges into the Barents Sea with a mean discharge of 197 m^3^ s^−1^ (years 2000 – 2020). Only 0.6% of the Teno River catchment is being used for human activity, making it one of the largest nearly undisturbed river ecosystems in Northern Europe. The Teno River’s drainage basin sits on ancient Precambrian rock, primarily composed of granite, granulite, gneiss, and schist. The soils in this area are diverse, ranging from glacial deposits to peatlands and residual soils. Our sampling sites were located on transects spanning from the shoreline up to the main riverbank, or across a sandbank between the main river and a small river-arm. On the main riverbank, the vegetation varied from graminoids (mainly *Carex* spp.) near the shore to evergreen dwarf shrubs (*Empetrum nigrum*) and up to 3-m-high deciduous shrubs (mainly *Salix* spp.) higher up on the riverbank. Mosses were important in the ground layer. The sandbank is mostly bare because of water erosion, but it has some vegetated patches, composed of *Carex* spp., *Salix* spp., and mosses. No organic layer was present in bare soils, whereas an organic layer of 0.5 to 5 cm depth had formed in vegetated soils. In the vegetated part of the main riverbank, the soils were classified as Fluvic Gleysol (Arenic, Skeletic) and on the sandbank, Stagnic Fluvic Gleysols (Arenic) and Fluvisol (Arenic) were found (IUSS Working Group WRB, 2022).

The Red River basin is situated in the discontinuous permafrost on the southern edge of Disko Island. It is much smaller than the Teno River catchment with an area of 100 km^2^, of which 72 % is land and 28 % is ice, and there is no human influence (Mankoff et al., 2020). The mean discharge of the Red River is 6.6 m^3^ s^−1^ (years 2015 – 2023). The soils in the region usually feature a shallow organic layer (around 5 cm) overlying mineral soil mixed with basaltic rock pieces (Peplau et al., 2025), but the riparian zone of the Red River is mostly bare of vegetation and has no organic layer because of wind and water erosion. In our sampling site, vegetation was present in small patches and composed of lichens and low-stature *Salix* spp., which had supported establishment of a 1 cm thick biocrust-layer (Ah horizon) underlain by a mineral horizon with a low organic content. In the bare soils, the mineral horizon at the surface is stratified with organic material due to wind erosion. The vegetated soils were classified as Fluvic Gleysol (Loamic, Skeletic) and bare soils as Fluvic Gleysol (Arenic, Skeletic) (IUSS Working Group WRB, 2022).

### 2.2 Soil sampling

The soil sampling at the two study sites was related to other on-going projects with different aims, which resulted in differences in the sampling scheme: At the Teno River, the spatial variability related to the distance from the shoreline was targeted, while at the Red River, the focus was on the differences between vegetated and bare soils, and between soil horizons. However, the sampling depth and the location in the direct vicinity of the river were similar at both sites, making the results well comparable. At the Teno River, soil samples were collected in the riparian zone in August 2023 from a total of 18 locations, 8 of which were bare and 10 vegetated (Figure 1b). The topsoil, 10 cm, including both the organic (when present) and mineral horizons, was sampled using a plastic soil corer with 5 cm diameter. Samples were collected along six transects, three of which (T1-T3) were in the riverbank of the main river channel, extending from the shoreline ∼30 m upslope. The other three transects (T4-T6) extended across a sandbank with a maximum width of 70 m and length of 220 m, located between the main river channel and a 10-12-m wide river arm. Samples were taken on each transect at three distances from the main shoreline (near, mid, and far).

At the Red River, soil samples were taken in September 2023 from four locations, two of which were bare and two vegetated. Soil samples were taken by a spade or by a steel corer with a diameter of 5 cm to a depth of ∼10 cm. Sampling locations were at an approximate distance of 2-4 m from the river, roughly corresponding to the mid distance at the Teno River (Figure 1c). Soil profiles were separated into layers according to visible differences in soil properties, making a distinction between organic (Ah) and mineral layers (C), which were further split according to differences in texture, redoximorphic features and/or stone content, resulting in a total number of samples of 19 (Figure S1). Pictures of the studied soils (Figure S2) and more details about the soil sampling of both sites can be found in the supplementary method section.

### 2.3 Soil properties

Soils from both sites were stored frozen for three months, thawed, and then stored at 5°C for two weeks prior to the experiments. Soil samples were cleaned of stones and roots and homogenized by sieving (2 mm mesh size). Soil properties including bulk density, WFPS, pH, electrical conductivity (EC), total C, total inorganic carbon (TIC), total N, extractable mineral N including NO_2_^−^, NO_3_^−^ and NH_4_^+^, DOC, DON, aromaticity of DOC, anions including chloride (Cl^−^), sulphate (SO_4_^2−^) and phosphate (PO_3_^−^), as well as microbial biomass were analyzed by standard methods in the laboratories of University of Eastern Finland and University of Hamburg. A detailed description of these methods is provided in the Supplementary Methods.

### 2.4 Soil N transformation rates

#### 2.4.1 Net mineralization and nitrification

For net nitrification and ammonification, 20 g of fresh soil was weighed in 120 ml-borosilicate bottles. The bottleneck was closed with pierced parafilm, and soils were incubated at +5°C for 14 days. At 5 time points (days 0, 1, 3, 7 and 14), an aliquot of 4 g of soil was taken and extracted with 16 ml of 1M KCl for determination of mineral N content. Concentrations of NO_3_^−^ and NH_4_^+^ in the soil extracts were determined by spectrophotometry as described in the Supplementary method. Net rates were calculated as the difference in mineral N content between the end and start of the incubation (0 to 14 days) divided by the incubation time. Net mineralization was calculated as the sum of net nitrification and net ammonification.

#### 2.4.2 Gross mineralization and nitrification

Gross mineralization and nitrification rates were determined in aerobic conditions at +5°C and field moisture content using the ^15^N pool dilution method (Kirkham & Bartholomew, 1955). In this method, the product pool (NH_4_^+^ for mineralization and NO_3_^−^ for nitrification) is isotopically enriched with ^15^N, and the process rate is derived from the dilution of this label over time.

In brief, 15 g of fresh soil was weighed into three parallel 50 ml centrifuge tubes (one per each sampling point) and dried on ice with table fans to reduce the water content so that after adding the labelling solution, the field moisture content was reached. In the mineralization experiment, we used for ^15^N labelling a solution of 10 at% ^15^(NH_4_)_2_SO_4_ (volume of 1.8 ml with 1.43 µmol N to 15 g FW, equal to 1.76 µg N g^-1^ DW). In the nitrification experiment, the label was 3 at% ^15^N-KNO_3_ (volume of 0.6 ml with 0.29 µmol N to 5 g FW, equal to 1.07 µg N g^-1^ DW). We used a stepper pipette to distribute the label evenly across the sample and then mixed using a glass rod. Samples were extracted at 3, 24, and 72 hours using 1M KCl.

The extracts were analyzed for NH_4_^+^ concentration (in mineralization experiment) and NO_3_^−^ (in nitrification experiment) as described in the Supplementary methods. For measuring the ^15^N at% in NH_4_^+^, we captured NH_4_^+^ into the solid form using the microdiffusion method (Lachouani et al., 2010) followed by analysis on EA-IRMS (Thermo Finnigan DELTA XP Plus IRMS, Flash EA 1112 Series Elemental Analyzer, and a Conflow III open split interface; both Thermo Finnigan). For measuring the ^15^N at% in NO_3_^−^, we used either the Titanium (III) method where NO_3_^−^ is chemically reduced to N_2_O (Altabet et al., 2019) or the denitrifier method which is based on bacterial conversion of NO_3_^−^ to N_2_O by *Pseudomonas aureofaciens* (Sigman et al., 2001; Casciotti et al., 2002), followed by ^15^N-N_2_O analysis on IRMS (Thermo Finnigan DELTA XP Plus IRMS) coupled with a gas bench for trace gas preconcentration and gas chromatography separation (PreCon-Gas Bench II; both Thermo Finnigan). A detailed description of the analytical methods can be found in the Supplementary methods. The gross rates were calculated between each two timepoints according to Hart et al. (1994). For gross mineralization, we report the initial rate between 3-24 hours. For gross nitrification, we report the rate between 24-72 hours, because before 24h we commonly observed an increase in the ^15^N label, violating the assumptions of the pool dilution method.

#### 2.4.3 Potential DNRA and denitrification

Potential DNRA rate was measured under anaerobic conditions (O_2_ < 0.1%) at +5°C in a soil slurry incubation with added NO_3_^−^ (Silver et al., 2001; Huygens et al., 2008). To minimize exposure to oxygen during pre-processing, soils were homogenized by hand in a glove bag under a pure N_2_ atmosphere. We weighed 25 g of fresh soil into four parallel 50 ml centrifuge tubes (one per each sampling point) and added 8 ml of solution containing 0.45 mmol N l^−1^ of 5 at% K^15^NO_3_ and 0.40 mmol N l^−1^ unlabeled NH_4_NO_3._ The mineral N addition was adjusted to ensure sufficient content of both mineral N species for ^15^N determination, equaling to 2.65 µg N-NO_3_^−^ g^-1^ DW and 2.36 µg N-NH_4_^+^g^-1^ DW. The labelling solution was purged with N_2_ prior to use to remove oxygen. During the 72-h incubation, samples were extracted at 3, 24, 48 and 72 hours using 1M KCl. The extracts were analyzed for NO_3_^−^ and NH_4_^+^ concentration and ^15^N at% as described above in section 2.4.2 and in Supplementary methods. DNRA rates were calculated between each two timepoints according to Huygens et al. (2008), using ^15^N enrichment in NH_4_^+^ and NO_3_^−^, mean NH_4_^+^ concentration in the time interval, and mean residence time of NH_4_^+^. Mean residence time was calculated by dividing the initial concentration of NH_4_^+^ by the gross consumption rate. Gross consumption was calculated by subtracting the net mineralization rate from the gross mineralization rate (see section 2.4.2). Here, we report the potential DNRA rate between 3-72 hours.

In the same experiment, we additionally determined the maximum rate of N_2_O production in the presence of added NO_3_^−^, later referred to as the potential denitrification rate. For this purpose, we took gas samples from the headspace at each time point before the soil extraction and analyzed the concentrations of N_2_O and calculated the maximum N_2_O production rate as described below in section 2.5. Importantly, these assays reflect the maximum microbial potential under optimized laboratory conditions and are not intended to represent actual *in-situ* field rates.

### 2.5 Greenhouse gas fluxes and production of N_2_O, CH_4_ and CO_2_

#### 2.5.1 *In-situ* GHG fluxes

*In-situ* fluxes of N_2_O, CH_4_ and CO_2_ from the soil surface were measured at the Teno River between September 09-16, 2024, and at the Red River between August 31-September 10, 2023. Fluxes were measured next to the soil sampling locations. Each plot was measured 6 times at the Teno River and 10 times at the Red River by a manual chamber technique using two portable greenhouse gas analyzers (LI-7820 for N_2_O/H_2_O and LI-7810 for CH_4_/CO_2_/H_2_O, LI-COR Biosciences, Lincoln, Nebraska, USA). A transparent cylindrical chamber was placed on permanently installed collars for measurement of net ecosystem CO_2_ exchange (NEE). Another opaque chamber measurement was done after covering the chamber with a reflective tarp and used for the determination of N_2_O, CH_4_ flux and ecosystem respiration (ER) in the dark. Fluxes were calculated based on the concentration change in the chamber headspace over time with linear regression fitting in MATLAB version R2023b using a published chamber flux script (Eckhardt & Kutzbach, 2016). Additionally, we measured N_2_O, CH_4_ and CO_2_ fluxes from the Teno River surface (n = 10) using manual floating chambers in August 01-03, 2023 (Saarela et al., 2025). Five syringe samples were taken during a chamber closure time of 21 min, and analyzed with a gas chromatograph (Agilent 7890B, USA). Detailed descriptions of soil and river flux measurements are given in Supplementary Methods.

#### 2.5.2 GHG production in soil incubations

The production rates of N_2_O, CO_2_ and CH_4_ in aerobic and anaerobic conditions were determined in incubation experiments at +5°C and field moisture content. In short, for anaerobic incubations, 15 g FW of fresh soil was weighed into sterile 120 ml glass flasks, flushed 4-5 times with pure N_2_ inside a glove bag (Kuopio; O_2_ < 0.05%) or in an anaerobic cabinet (Hamburg; O_2_ < 0.006%) and sealed with rubber septa and screw caps. For aerobic treatment, laboratory air was used for headspace. During the incubation of 7 days, gas samples were taken from the incubation bottles on 6-8 occasions and analyzed on a gas chromatograph (in Kuopio Agilent 7890 A, in Hamburg Agilent 7890 B). Gas production rates between time points were calculated from concentration changes (ppm day^-1^) using the ideal gas law. Based on visual inspection of the CH_4_ and CO_2_ concentrations, we identified the period of stable gas production or consumption rate as 3 days and fitted a linear regression to the measurement points during that period. For N_2_O, we report the maximum production between any two time points for both anaerobic and aerobic treatments, because we sometimes observed initial production followed by N_2_O consumption or stable concentration, particularly in the anaerobic treatment. Gas production rates were normalized to soil dry weight.

### 2.6 Statistical analysis

The plotting and statistical analysis were done on R (RStudio, version 3.6, Posit Software, PBC (RStudio Team, 2022). The differences between vegetated and bare soils in soil properties and N transformation rates at both sites and *in-situ* GHG fluxes at the Red River were tested using T-test (*stats* package). For the Teno River, we used two-way ANOVA (*stats* package) to test differences in N and C pools, i*n-situ* GHG fluxes, GHG production rates and N transformation rates according to two factors: 1) distance from the main shoreline and 2) between bare and vegetated sites, as well as their interaction. Prior to statistical tests, the normal distribution of the data was tested using the Shapiro-Wilk test and visual inspection of histograms. Right-skewed data were log-transformed to achieve normal distribution. Principal component analysis (PCA) including all variables (except for those with several missing values, or no variability) was done separately for each site with ‘*factoextra’* package (Kassambara & Mundt, 2020), with the aim to explore the differences between bare and vegetated soils and different soil layers, as well as the importance of different variables for data clustering. Data for PCA were scaled and centralized. With the same dataset, Pearson correlation matrix was done with *corrplot* package (Wei & Simko, 2024). For correlation analysis, variables were log-transformed when needed to achieve a normal distribution. The dominance of N pools and turnover processes in the bare and vegetated soil of each study site was visualized with Sankey diagrams using ‘*ggalluvial*’ package.

## 3. Results

### 3.1 Soil properties including N pools

Riparian soils at both sites exhibited low-to-intermediate C content and C/N ratios (Table S1, Figure 2). The presence or absence of vegetation had a significant effect on many soil traits: At both study sites, vegetated soils were wetter and more acidic than bare soils, and had a higher total N content, C content, C/N ratios, and electrical conductivity (EC). Vegetated soils also contained more fine soil particles (silt, clay). Extractable N and C pools were generally higher in vegetated soils, especially at the Teno River. At Teno, DON dominated the extractable N pool in vegetated soils, whereas NH₄⁺ was dominant in bare soils; at the Red River, DON comprised most of the extractable N forms in both bare and vegetated soils (Figure 3). From mineral N forms, NO_3_^−^ was dominant at the Red River in both bare and vegetated soils, while the Teno River had more NH₄⁺. Total C was consistently very low in bare soils (Teno River: 0.1%; Red River: 0.4%). In vegetated soils, total C was similar at both study sites, but total N was nearly twice as high at the Teno River, resulting in a lower C/N ratio (Teno River: 9.6; Red River: 18.7 (w/w)). Lower pH of 5.1–5.9 was found at Teno River than at the Red River, where it was 6.7–7.1. A detailed comparison of the biogeochemical and physical properties of the Teno River and the Red River riparian zones is provided in Supplementary Table 1.

**Figure 2.**
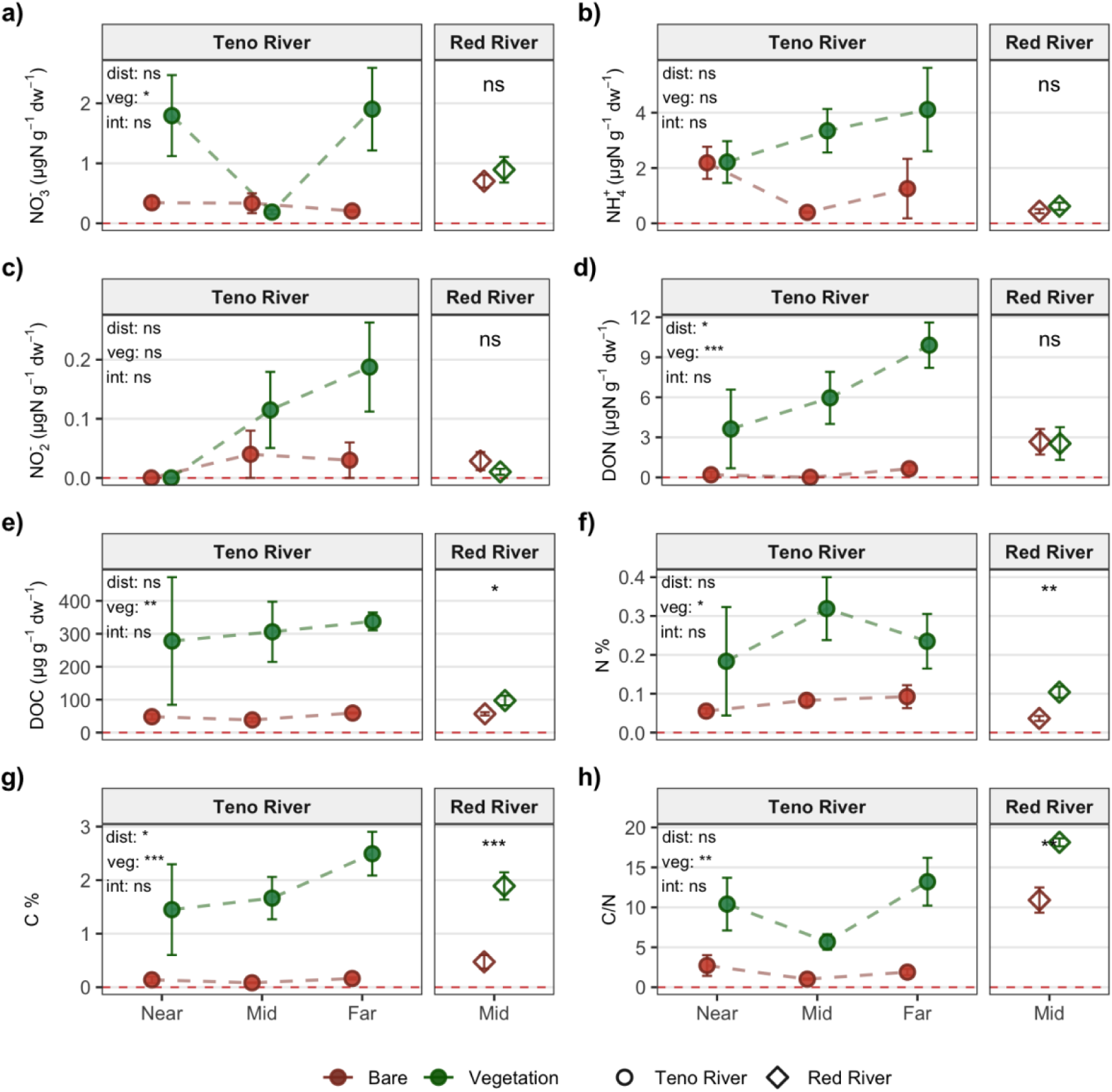
Effect of distance from the shoreline and vegetation on N and C content: mineral N species, including NH, NO_2_^−^, and NO_3_^−^ **(a, b, c)**, DON **(d)**, DOC **(e)**, total N **(f)**, total C **(g)**, and C/N ratio **(h)** in bare (brown) and vegetated (green) soils. At the Red River, data represent mean values across soil layers. Standard deviations are shown as error bars. Significant differences between bare and vegetated soils according to the T-test are shown for the Red River. For the Teno River, significant differences according to a two-way ANOVA are shown for distance (d), vegetation cover (v) and their interaction (int) by *** = *p* < 0.001, ** = *p* < 0.01, *= *p* < 0.05, ns = not significant.

**Figure 3.**
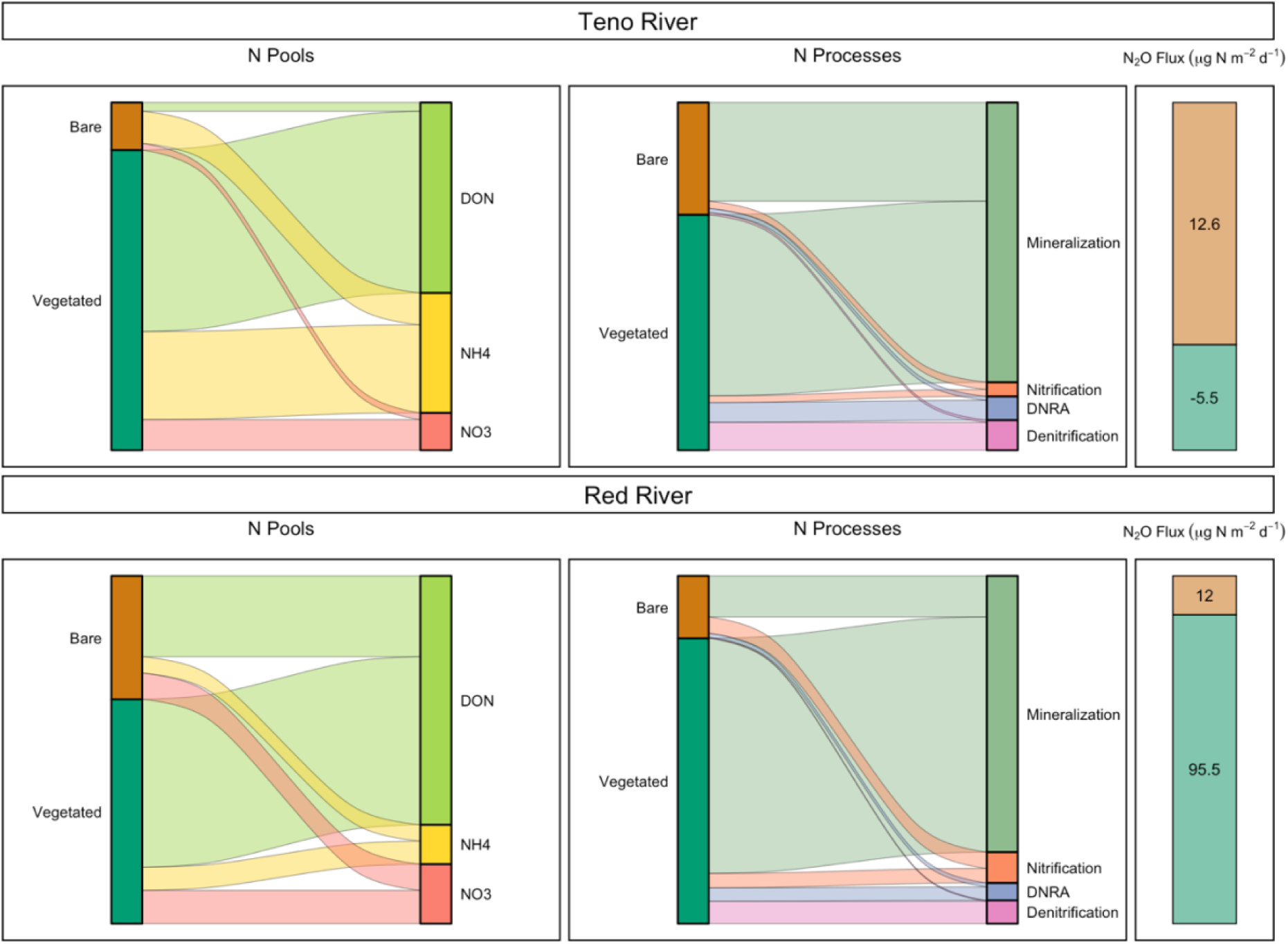
Dominance of N pools and transformation processes in riparian soils at the Teno River and the Red River. Sankey diagrams illustrate the relative importance of extractable N species **(left)** and N transformation processes **(mid)** and N_2_O emissions **(right)** in bare and vegetated soils and are based on mean values. Although trace amounts of NO_2_^−^ were found, it is not included in the diagram because of the low contribution.

**Figure 4.**
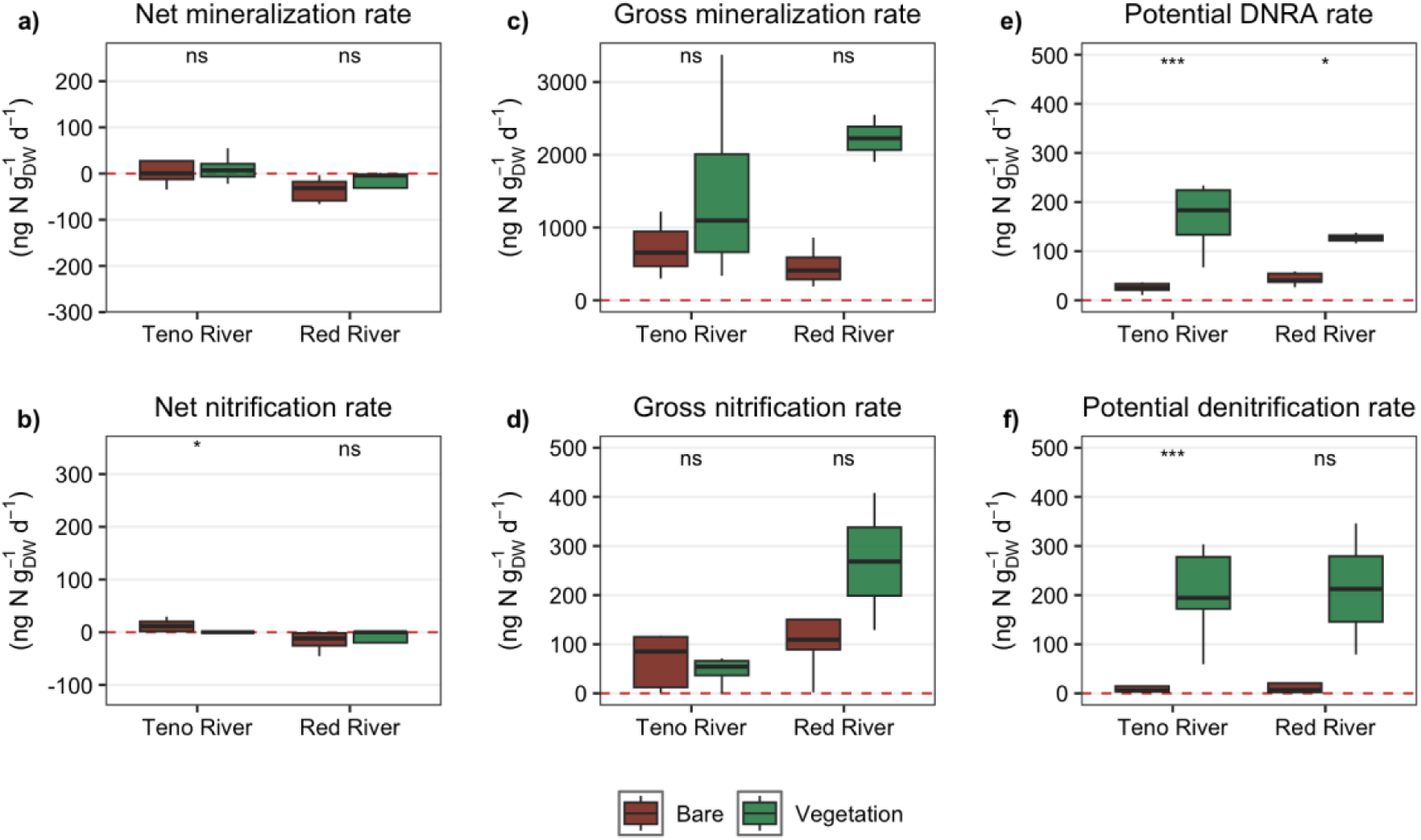
Comparison of N transformation rates between the Teno River and the Red River and between bare (brown) and vegetated (green) soils: net mineralization and nitrification **(a, b)**, gross mineralization and nitrification **(c, d)**, and potential DNRA and potential denitrification **(e, f)**. Box plots show lower and upper quartiles, median (thick black line), smallest and largest values (thin black line), and outliers (circles). T-test significance is by *** = *p* < 0.0005, ** = *p* < 0.005, * = *p* < 0.05, ns = not significant.

The different sampling schemes at the two sites allow us to explore different spatial patterns in soil properties. At the Teno River, many soil properties including C and N pools showed significant differences according to the distance from the shoreline (Figure 2). In vegetated soils, NO_2_^−^, DON, and total C content nearly doubled with increasing distance (Figure 2 c, d, g). This was associated with an increase in acidity and silt and clay content and a decrease in bulk density and sand content (Figure S3). The trend for NO_3_^−^ was different, with the lowest values occurring at mid distance, and a relatively high NO_3_^−^ at near distance, similar to NH₄⁺ (Figure 2 a, b). Overall, bare soils showed less variability and weaker distance-related trends for C and N pools compared to vegetated soils. Soil samples at the Red River, collected from mid distance and separated into different soil horizons, exhibited distinct depth patterns for some soil properties (Figure S4). In vegetated soils, total C, N, DOC, and DON were higher in the surface biocrust than in the mineral soil layer. In contrast, in bare soils these pools increased with depth. In vegetated soils, C/N ratios in the biocrust and mineral horizon were similar, while bare soils showed depth variability from 3 at the surface to 20 at ∼10 cm. Fine particle and moisture content increased with depth in bare soils but remained stable across depths in vegetated soils (Figure S3).

### 3.2 Soil N transformation rates

Nitrogen transformation rates were generally similar in magnitude at the Teno River and the Red River, but the presence of vegetation substantially increased the mineral N turnover at both study sites (Figure 3c-d, 4, 5). This difference between vegetated and bare soils was particularly remarkable for potential denitrification rate, with a 36-fold difference in median value at the Teno River (p = 0.0001) and a 32-fold difference (albeit not significant) at the Red River (p = 0.4). Similarly, DNRA rates were significantly higher in vegetated soils at both sites (Teno: p = 0.0001; Red River: p = 0.03), with an 8-fold and 4-fold difference at the Teno River and the Red Rivers, respectively. Gross mineralization also followed this pattern, with a tendency towards higher rates in vegetated than bare soils at both study sites (Teno River:1094.6 vs. 655.3 ng N gDW^−1^ d^−1^ and Red River: 2225.8 vs. 434.3 ng N gDW^−1^ d^−1^). Further “downstream” mineral N turnover processes nitrification, denitrification, and DNRA were low compared to gross mineralization, even though denitrification and DNRA were measured with added NO_3_^−^ and therefore represent potential rates (Figure 3c-d, 4). Gross nitrification rates at the Teno River were similar in bare and vegetated soils, representing a notable exception to the general pattern of higher rates in vegetated soils. From the two alternative NO_3_ ^−^ reduction pathways, vegetated soils showed similar potential rates of DNRA and denitrification, while in bare soils the DNRA potential was higher. Net mineralization was predominantly negative at the Red River and close to zero at the Teno River, with a few positive rates. Net nitrification was consistently positive in Teno bare soils, but negligible or negative elsewhere.

At the Teno River, distance from the shoreline did not significantly influence N transformation rates, except for potential DNRA (p = 0.04), which peaked at both near and far distances, as did NO_3_^−^ (Figure 5f). Denitrification showed a roughly opposite pattern, peaking at mid distance (Figure 5e). Gross mineralization was clearly higher in vegetated than in bare soils at mid distance, but independent of the presence or absence of vegetation at other distances. Gross nitrification was markedly higher in bare than vegetated soils at near distance, but at mid and far distances the rates were similar between vegetation categories. At the Red River, there were no clear depth patterns in mineralization nitrification, or denitrification rates, which often varied more between sampling locations than between soil layers. In vegetated soils, some rates (nitrification, denitrification) tended to be higher in the biocrust than the mineral layer underneath. Bare soils showed slight increases of net nitrification, gross mineralization and potential denitrification with depth (Figure S5).

**Figure 5.**
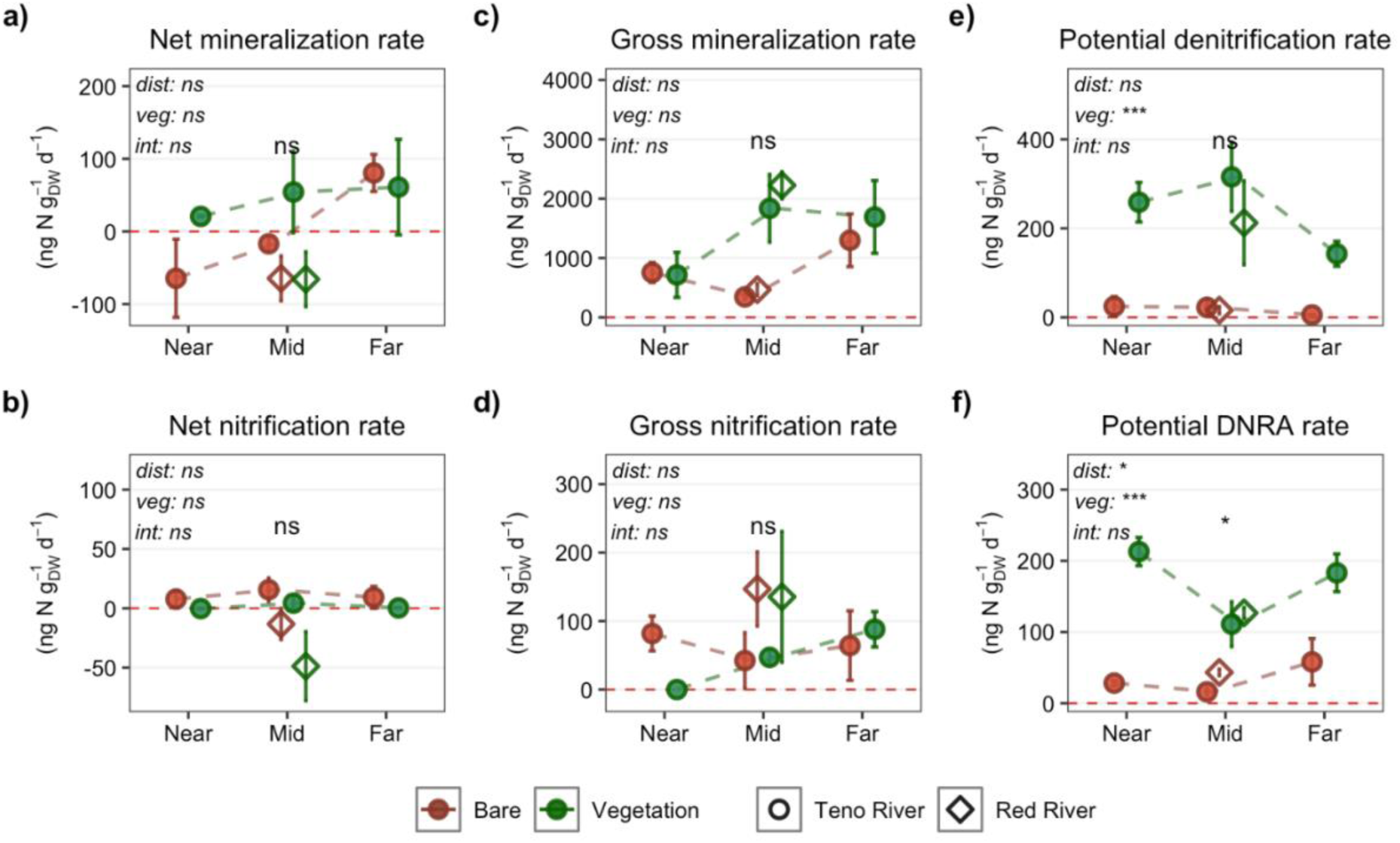
Effect of distance from the shoreline and vegetation on nitrogen transformation rates, including net mineralization and nitrification **(a, b)**, gross mineralization and nitrification **(c, d)**, as well as potential denitrification and potential DNRA **(e, f)** at the Teno River and the Red River in bare (brown) and vegetated (green) soils. At the Red River, data represent mean values across soil layers. Standard deviations are shown as error bars. Significant differences between bare and vegetated soils according to the T-test are shown for the Red River. For the Teno River, significant differences according to a two-way ANOVA are shown for distance (d), vegetation cover (v) and their interaction (int) by *** = *p* < 0.001, ** = *p* < 0.01, * = *p* < 0.05, ns = not significant.

### 3.3 Greenhouse gas dynamics

Riparian *in-situ* N_2_O fluxes at the Teno River were generally low, ranging from small sinks to small sources (–78 to 119 µg N m^−2^ d^−1^; Figure 6). The river surface showed higher N_2_O emissions than the riparian soils (127.1 ± 13 µg N m^−2^ d^−1^). A significant difference was found at Teno between bare soils, which were N_2_O sources, and vegetated soils, which acted as N_2_O sinks. Riparian soil N_2_O emissions increased with distance from the shore: sources from bare soils intensified, and sinks in vegetated soils weakened. At the Red River, *in-situ* N_2_O fluxes ranged from small sinks in bare soils to moderate sources in vegetated soils (–63 to 384 µg N m^−2^ d^−1^; Figure 6). In contrast to the flux pattern observed at the Teno River, significantly higher N_2_O emissions were found at the Red River in vegetated than in bare soils.

**Figure 6.**
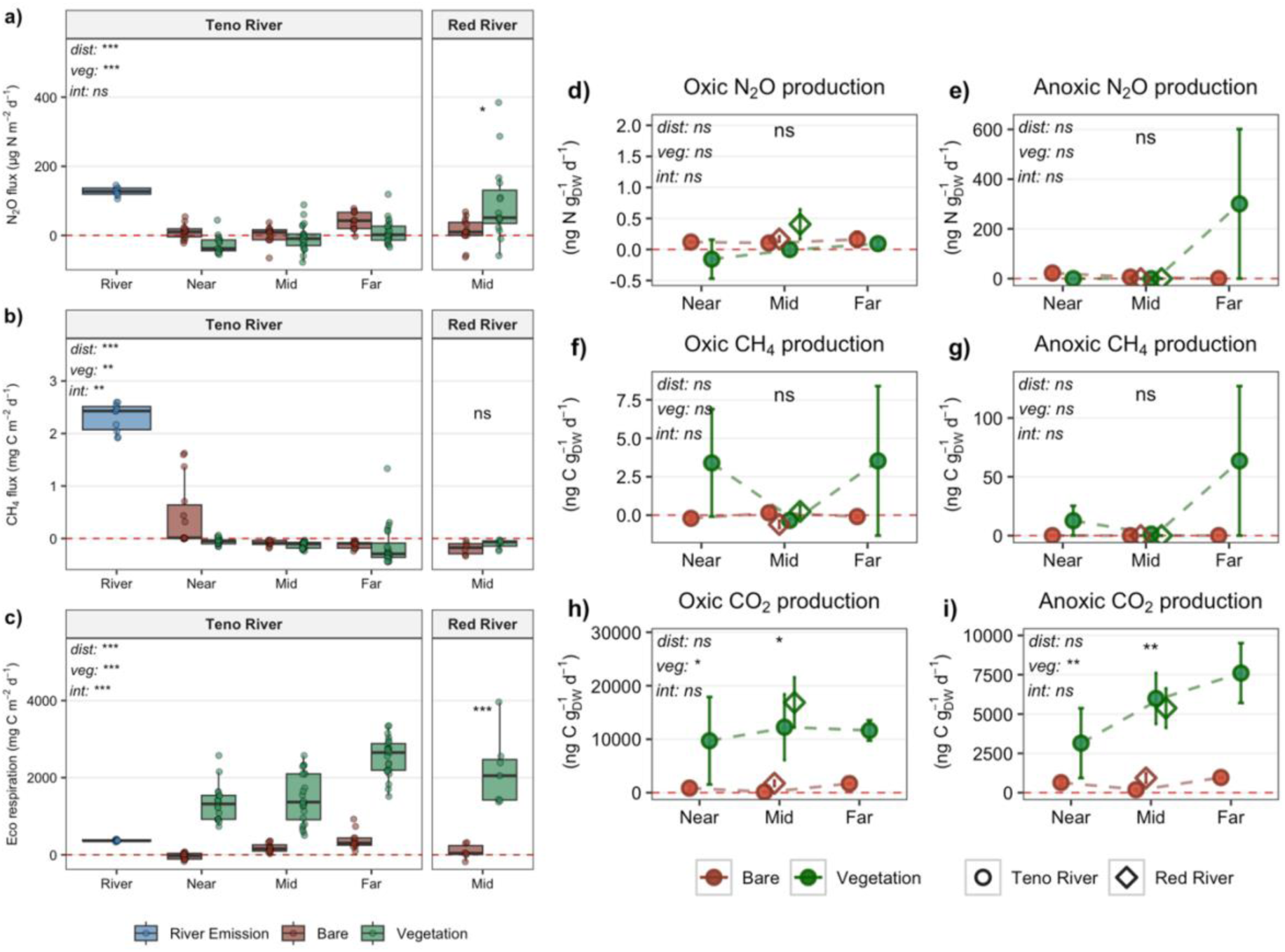
Greenhouse gas dynamics of riparian soils at the Teno River and the Red River: I*n-situ* fluxes of N_2_O **(a)**, CH_4_ (**b)** and CO_2_ (**c)** and GHG production in oxic (**d, f, h)** and anoxic incubations **(e, g, i)**. For the Teno River, also *in-situ* emissions from the river surface are shown. Different colors indicate emission from the river surface (blue), bare soils (brown) and vegetated soils (green) along the river-to-upland gradient. Box plots show lower and upper quartiles, median (thick black line), smallest and largest values without outliers (thin black line), and the raw data (circles). Point plots **(d to i)** show mean values with standard deviations for both sites. Significant differences between bare and vegetated soils according to the T-test are shown for the Red River. For the Teno River, significant differences according to a two-way ANOVA are shown for distance (d), vegetation cover (v) and their interaction (int) by *** = *p* < 0.001, ** = *p* < 0.01, * = *p* < 0.05, ns = not significant.

For CH_4_, the riparian zone at the Teno River acted as a net *in-situ* sink (–0.02 ± 0.3 mg C m^−2^ d^−1^), with significantly stronger uptake in vegetated soils and with increasing distance from the shoreline (Figure 6). Net CH_4_ emissions were found in bare soils near the main shoreline and in vegetated sandbank soils next to the river’s arm. Similar to N_2_O, the river was a CH_4_ source (2.3 ± 0.2 mg C m^−2^ d^−1^), with emissions exceeding those from riparian soil. At the Red River, both bare and vegetated soils acted as small net CH_4_ sinks. Ecosystem respiration (ER) was significantly higher in vegetated soils at both study sites and increased with distance from the shoreline at the Teno River (Figure 6). Bare soil ER at far distance (next to river arm) reached rates comparable to river emissions. The spatial pattern of ER reflected the spatial pattern in gross primary production (GPP) (Figure S6). Notably, bare soils exhibited CO_2_ uptake (both GPP and NEE) at mid and far distances despite the absence of plants. At the Red River, ER was 50-fold higher in vegetated than bare soils. The vegetated soils also showed high net ecosystem exchange (NEE), while bare soils did not exhibit net CO_2_ uptake under light conditions, in contrast to the Teno river (Figure S7).

Oxic incubations with the Teno River soils mirrored the *in-situ* N_2_O patterns, with bare soils producing more N_2_O than vegetated soils, although not significantly so. At the Red River, oxic N_2_O production decreased with depth, particularly in vegetated soils, where biocrust showed higher N_2_O production than the underlying mineral horizon (Figure S8). Under anoxic conditions, no N_2_O production was detected except for one vegetated Teno sample. In contrast to net CH_4_ uptake observed *in situ,* vegetated soils at the Teno River showed net CH_4_ production in both oxic and anoxic incubations. At the Red River, the highest CH_4_ uptake was found in bare surface soil and the highest CH_4_ production in the biocrust of vegetated soil, while no CH_4_ production was detected under anoxic conditions. At both study sites, CO_2_ production was consistently higher under oxic than anoxic conditions and significantly higher in vegetated than bare soils. The spatial patterns of oxic and anoxic CO_2_ production roughly mirrored those in total C content with respect to both distance from the shoreline at the Teno River and depth at the Red River.

### 3.4 Drivers of spatial variability in N pools and processes

Principal component analysis (PCA) confirmed that soil properties at the Teno River including N and C pools, N transformation rates and GHG dynamics are strongly differentiated by the presence or absence of vegetation (Figure 7a). The first PCA axis explained 44% of the total variability in the data and clearly separated vegetated and bare soils based on organic matter (total C, DON, DOC), WFPS, and pH. The second axis explained 13% of the variability and was negatively related to net ammonification, net mineralization, and C/N ratios, and positively related to CH_4_ flux, clay content, and NH_4_^+^ content (Figure S9). Collectively, these components distinguished vegetated soils, characterized by higher C content, finer texture, and wetter conditions, all favoring potential denitrification and DNRA, from bare soils with higher nitrification rates and N_2_O fluxes. At the Red River, primary separation occurred between soil horizons rather than vegetation categories. The first axis (42% of the variability) distinguished organic-rich horizons (A, Ab, and CA) from organic-poor horizons (C), primarily driven by total C, CO_2_ production, DOC, EC, and pH (Figure 7b). The second axis (24% of the variability) was positively related to NH_4_^+^, NO_3_^−^, C/N ratio, and net mineralization, and negatively to DON and aerobic N_2_O production (Figure S9).

**Figure 7.**
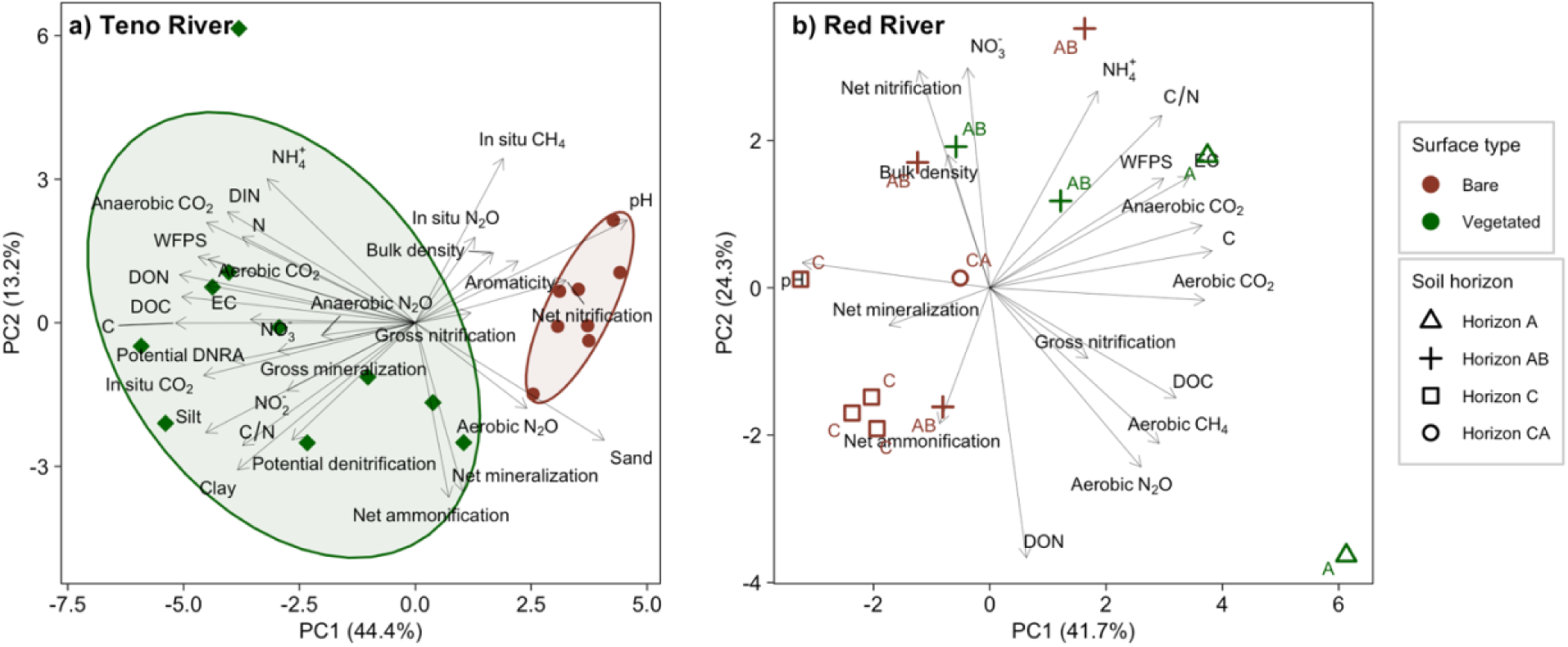
Biplot of the first two components of principle component analysis (PCA) including soil properties including N pools, N transformation rates, as well as greenhouse gas fluxes and production rates and at the Teno River **(a)** and the Red River **(b)**. In **a)**, bare and vegetated soils are shown with brown and green symbols and ellipses (confidence level 0.8), respectively. In **b)**, similar color codes are used for bare and vegetated soils, while soil horizons A, Ab, CA and C are indicated by different shapes.

**Figure 8.**
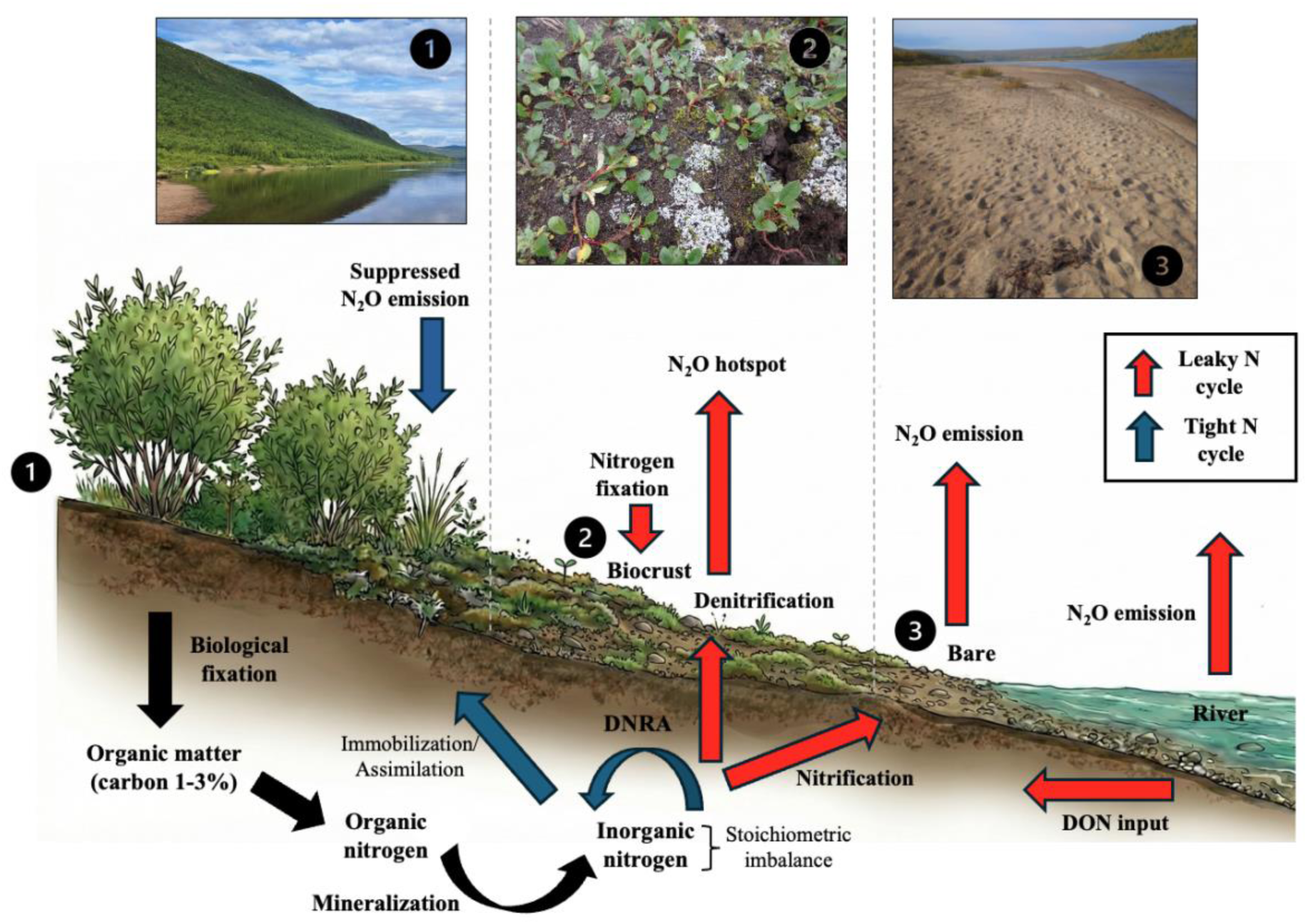
The conceptual figure illustrating the effect of vegetation type and substrate stoichiometry on N cycle in Arctic riparian zones and the dominance of retention vs. loss processes. **(1)** Dense vascular vegetation cover at the Teno River creates a tight N cycle, indicated here by blue arrows. Vascular plants improve C availability via organic matter inputs and root exudation, which enhances organic matter accumulation and N turnover, but at the same time they compete for mineral N. This promotes N retention processes, including DNRA, effectively suppressing N_2_O losses. **(2)** Biocrusts at the Red River, composed of lichens with cyanobacterial associates, act as ecosystem engineers enhancing soil N and C, thereby creating hotspots for N_2_O emissions, likely due to the coupled nitrification-denitrification process. **(3)** Bare soils at Teno sandbank show active N mineralization and turnover, resulting in a leaky N cycle, indicated here with red arrows. We propose that mineral N cycling in these soils with very low C content is fueled by DON infiltration from the river. C-limitation, as indicated by low C:N ratio, forces the microbiome to mineralize organic N for energy, resulting in high inorganic N availability and N_2_O emissions. The variability in vegetation cover in Arctic riparian zones from bare soil through biocrust to dense vascular plant cover defines the dominance of N loss versu retention processes.

Pearson correlation analysis further supported these findings, highlighting that soil moisture and fine material were consistently higher in vegetated soils and positively correlated with C and N pools (Figure S10, S11). At Teno, anaerobic pathways denitrification and DNRA were positively associated with each other and WFPS and showed negative correlation with net nitrification. The positive correlation between these reductive processes with C content and ER at both study sites points towards C limitation. At the Teno River, gross mineralization correlated positively with total C (R = 0.56, p = 0.0005) and total N (R = 0.36, p = 0.01). *In-situ* N_2_O flux at Teno was negatively correlated with potential denitrification (R = 0.25, p = 0.03) and C/N ratios (R = 0.23, p = 0.04).

## 4. Discussion

### 4.1. Arctic riparian zones are sites of active mineral nitrogen cycling because of substrate stoichiometry

Although riparian zones in general are described as the “kidneys” of river systems because they reduce nutrient inputs from land to streams, this filtering capacity is thought to be low in boreal and Arctic systems where biogeochemical processing and nutrient retention are limited by low temperatures (Pinay et al., 2018). With this in view, our soils exhibited surprisingly high N filtering capacity at the low incubation temperature, primarily driven by retention (mineralization, immobilization, DNRA) rather than loss pathways (nitrification, denitrification). Our study revealed that high-latitude riparian soils show relatively high rate of gross N mineralization. Thereby, they reduce the lateral N inputs to the river by converting DON to NH_4_^+^, which is less mobile and more easily assimilated by plants and microbes. A comparison with Pan-Arctic synthesis data reveals that our soils exhibited notably higher gross mineralization rates than predicted by C content (Ramm et al., 2022, Figure S12), especially the bare ones. This can be explained by the extremely low C/N ratios, similar or even lower than the stoichiometric ratio in microbial biomass (Cleveland & Liptzin, 2007; Smith et al., 2024), as previously observed in other C poor soils (Agnelli et al., 2021; Matschullat et al., 2018; Mörchen et al., 2019). The low C/N ratios suggest high N availability relative to C. In such C limited environments, the microbial community utilizes organic N compounds as energy and carbon sources rather than solely for N acquisition (Pessina et al., 2025, Elrys et al., 2021a, 2021b). The low net mineralization rates (Figure 4), however, indicate rapid re-immobilization of mineralized NH_4_^+^, contributing to overall N retention in the riparian zone. This rapid immobilization reduces the availability of substrate for nitrification, which represented only 10% of total mineralized N (Figure 3). The fraction of gross nitrification from gross mineralization was lower than in the Pan-Arctic synthesis data (Ramm et al., 2022).

Importantly, we were also able to detect the rarely studied DNRA pathway in almost all samples, further demonstrating N retention potential of the Arctic riparian soils. Potential DNRA rate was similar to potential denitrification rate in vegetated soils, and in the bare soils, it was even the dominant NO_3_^−^ reduction pathway, supporting the previously published genetic evidence (Sagai et al., 2023). This is in contrast with the literature where denitrification is often highlighted as the dominant NO_3_^−^ consumption pathway (Pinay et al., 2018), and the possibility of DNRA is neglected. In our study, DNRA increased with soil C content and was higher in vegetated soils with very high DOC/NO_3_^−^ ratios (Figure 2), in accordance with the literature (Ye et al., 2023). It must be noted that these potential rates under strict anoxia and with NO_3_^−^ supplements may not accurately mimic the real in situ conditions, but still, they provide valuable insights to the potential relative importance of DNRA. Future research should further investigate DNRA under variable redox conditions and substrate availability to better constrain the importance of this pathway in Arctic riparian soils.

### 4.2. Vegetation regulates Arctic riparian nitrogen cycling through carbon supply and soil stabilization

The presence of vegetation strongly shapes soil biogeochemical cycling (Figure 7); yet its role for soil N turnover is complex, as it can both enhance and reduce mineral N availability for soil microbes. At the Teno River, the 20-fold greater C content in vegetated soils compared to bare soils was clearly driving the elevated N turnover rates. Vegetation contributes to the SOM pool by litter inputs but also protects the soils against erosion by wind and flowing water, supporting the accumulation of SOM and fine particle to accumulate and the formation of biocrust (Figueiredo et al., 2016). Given the relatively low C content of the studied riparian zones, labile C inputs via root exudation and plant litter may enhance N turnover, particularly N mineralization (N mining; Bengtson et al., 2012; Wild et al., 2016, 2017) and the heterotrophic denitrification process (Booth et al., 2005; Burgin & Hamilton, 2007; Chapuis-Lardy et al., 2007). On the other hand, plants compete with microbes for available N, thereby limiting the NH_4_^+^ supply for nitrification and NO_3_^−^ supply for denitrification. Therefore, Arctic N_2_O hotspots are commonly found in unvegetated or sparsely vegetated areas (Voigt et al. 2020, Marushchak et al., 2011), leading to our hypothesis of higher N losses from bare riparian soils. While our results altogether suggest that N losses from Arctic riparian zones are rather minor, at the Teno river we found some evidence for a leaky N cycle in bare soils. Although we observed significantly higher extractable N pools in vegetated soils, bare soils had higher N_2_O emissions. An interesting finding is that the opposite was true for the Red River: There, *in-situ* N_2_O emissions were higher in vegetated than in unvegetated soils and reached a maximum flux of 384 µg N m⁻^2^ d⁻^1^, which is an order of magnitude higher than the median for permafrost-affected soils (Voigt et al. 2020).

We identify two main mechanisms that may explain the opposite effect of vegetation on N_2_O emissions at the two study sites: Firstly, at the Red River vascular vegetation was sparse and low-statured and, consequently, the plant N uptake was lower than in vegetated soils at the Teno River (Figure 1). Secondly, while the bare soils at the Red River did not have any organic layer at the surface, a well-developed biocrust was found in the vegetated patches, presumably due to physical protection against erosion by vascular plants. The role of biocrusts as ecosystem engineers is well-described in other polar environments: Biocrusts not only stabilize the soil surface and accumulate fine sediments but can also facilitate biological N_2_ fixation via cyanobacterial associates (Belnap, Weber & Büdel, 2016; Stewart et al., 2014; Fulweiler et al., 2025; Tschitschko et al., 2024). By coupled inputs of N and labile C, biocrusts may fuel microbial activities, including coupled nitrification-denitrification. Indeed, vegetated soils at the Red River exhibited the highest gross nitrification rates among the studied riparian soils, and both gross nitrification and aerobic N_2_O production were higher in the biocrust layer than the underlying soil (Figure S8). Taken together, our results suggest a complex role of vegetation in regulating soil N cycling, which is important to consider when assessing the impacts of Arctic greening and shrubification (Mekonnen et al., 2021; Zhou et al., 2019), as well as vegetation changes related to thermokarst disturbances (Zheng et al., 2025), and glaciers retreat (Agnelli et al., 2021).

### 4.3. Riverine inputs fuelled nitrogen turnover in shoreline soils

While vegetation was the main factor controlling biogeochemical cycling in the studied riparian zones and largely explained the changes along the distance gradient from the Teno River shoreline, we also observed significant spatial heterogeneity that did not follow this pattern. The relatively high gross mineralization and nitrification rates in the bare sandbank, despite the low SOC content, suggest that these processes may be fuelled by N inputs from the river. In these coarse-textured, permeable soils, continuous infiltration of riverine DON can be important in providing substrate for microbial activity. This aligns with the concept of biogeochemical hotspots at aquatic-terrestrial interfaces, where hydrologic mixing creates optimal conditions for active N and C biogeochemistry (Vidon et al., 2010). The high level of in-stream N cycling activity is implied by the higher N_2_O emissions from the river surface as compared to the riparian soils. Our interpretation is further supported by detection of N_2_O and CH_4_ emissions (up to 118.9 µg-N m⁻^2^ d⁻¹ and 1.5 mg C m⁻^2^ d⁻¹) in bare soils directly adjacent to the Teno River. Our results suggest that the dominant pathway of N2O release from the sandbank is nitrification, given the low moisture content of these sandy bare soils (WFPS < 55%) and the negative correlation between *in-situ* N_2_O emissions and denitrification potential (Figure S10).

### 4.4. Importance of N cycling in riparian zones in the changing Arctic

Our findings highlight the potential significance of riparian zones in regulating N export from land to river in the rapidly warming Arctic, where vast stores of soil organic C and nutrients are being mobilized. In this context, the poorly understood N biogeochemistry of the riparian soils represents a missing piece in understanding the effect of in-land processes on aquatic biogeochemistry. In addition to investigating changes in permafrost N release and plant N acquisition in the catchment area (Speir et al., 2024) and N cycling within rivers, elucidating N dynamics in riparian soils may help explain the recent increases in DON and decreases in NO_3_⁻ in Arctic rivers (Tank et al., 2023). Traditional assessments of riparian buffering capacity have in general focused on specific processes like denitrification, missing a holistic view of the N cycle (Pinay et al., 2018), and for high-latitude river system this information is even more scarce. By characterizing several important mineral N transformation processes, we addressed this gap and revealed that Arctic riparian zones are not inert with respect to N despite the low C and N content and cold temperatures; rather, they operate as active biogeochemical zones. A clear indication of this was the high mineralization efficiency which, depending on the location, supported either rapid microbial immobilization or N loss pathways. Importantly, high gross mineralization did always not translate into high gaseous N losses, at least in the form of N_2_O.

As the Arctic warms, hydrological conditions are anticipated to become more variable, potentially increasing the extent of disturbed, bare riparian zones (Bring et al., 2016; Lind et al., 2014). At the same time, warming is expected to promote shrub expansion in stable areas (Naito & Cairns, 2011; Michael et al., 2005). Our data suggest that this dual trajectory will have complex implications for the N cycle. Vegetation expansion may enhance N retention through plant uptake and DNRA, while increased river dynamism, bare soil and biocrust exposure could enhance nitrification or coupled nitrification-denitrification driven N_2_O losses. Comprehending this balance between the tight N cycle (vegetated) and the leaky N cycle (bare and biocrust) of riparian soils is crucial for predicting the future N dynamics of the changing Arctic. We propose further mechanistic studies and microbial investigations to confirm our findings and shed light on the role of processes not directly measured here, such as N fixation and N_2_O reduction to N_2_. Furthermore, the seasonality in riparian N cycling should be a key priority in future studies.

## 5. Conclusion

Expanding on earlier studies that typically examined isolated N processes, our comparative analysis of the N cycle in two pristine Arctic River riparian zones provides a more complete biogeochemical picture. It reveals a common pattern: despite distinct geographical settings, N cycling within these riparian zones is governed by severe organic matter limitation. We explain the paradox of high N turnover in C poor soils by identifying stoichiometric constraints as the primary driver. While vegetation was associated with higher mineral N turnover at both study sites, it had opposite effects on GHG emissions depending on its type and intensity. This distinction is vital for predictive modelling, as they suggest that the climate feedback of Arctic greening will depend heavily on whether succession favours vascular shrubs or cryptogamic covers. Finally, the consistent detection of potential DNRA across almost all samples highlights the importance of this pathway overlooked in riparian soils. DNRA appears to be a key retention mechanism at the terrestrial-aquatic interface, potentially reducing N export to aquatic systems. Collectively, our results reframe Arctic riparian zones as highly active biogeochemical zone. As the Arctic continues to warm, the interplay between hydrologic conditions in the soils and changes in vegetation coverage and composition may significantly modify the nutrient stoichiometry, with potential implications for GHG footprint of high-latitude riverine landscapes.

## Data availability

The data that support the findings of this study is openly available at https://doi.org/10.6084/m9.figshare.29976313.v2

## Acknowledgements

This work was funded by the Research Council of Finland in the frame of projects Thaw-N (no. 349503) and FinNcycle (no. 29310001771) Federal Ministry of Research, Technology and Space (BMFTR) DAAD PPP Finland exchange project 2023-2025 FinNcycle (project ID 57663850), C. F. was additionally supported by Federal Ministry of Research, Technology and Space (BMFTR) project MOMENT (no. 03F0931A) and the Cluster of Excellence ‘CLICCS-Climate Change, and Society’, contributing to the Center for Earth (CEN) University of Hamburg. L.K. received funding from the Research Council of Finland (MeMo2, no 354501) and holds a fellowship funded by the University of Eastern Finland [no grant number]. H.M.P.S received funding from the Research Council of Finland (NitroBiome, no 342362). Soil samples in Finland were collected during an expedition in August 2023 supported by the ACCC Flagship funded by the Research Council of Finland project number 357905. Soil samples in Greenland were collected during Expedition 08/09-2023 to Greenland with access granted by the non-exclusive licence no. G23-101 for utilization of Greenland genetic resources (Prior informed consent under the Biological Diversity Convention and the Nagoya Protocol) issued by the Ministry of Foreign Affairs, Business and Trade. We acknowledge conducting fieldwork on the ancestral land of the People of Kalaallit Nunaat, and honour with appreciation of the land itself and the Kalaallit People. We thank Birgit Grabellus, Selina Undeutsch, Johanna Kerttula, Lisa Beckers, Yvonne Wiese, Paula Martínez-Risco Martínez, Markus Ankele and Jan Penopp for their assistance during field and laboratory work, and Taija Saarela for help with river flux calculations.

