## Supplemental Files for "Soil nitrogen dynamics in riparian zones of two Arctic rivers"

*Journal of Soil Biology and Biochemistry*

Contents of this file

Supplementary methods

Figures S1 to S12

Table S1

### Supplementary methods

#### Study sites

Vegetation in the Teno River valley consists mainly of trees and shrubs (*Betula nana*, *Betula pubescens* subsp. czerepanovii, *Salix* sp., *Vaccinium*), mosses, and grasses (*Calamagrostis*, *Avenella*). The upper catchment area in the higher altitude area is a mixture of sub-arctic/arctic peatlands and fell vegetation. On the vegetated shoreline, the soils were classified as Fluvic Gleysol (Arenic, Skeletic). On the sandbank, Stagnic Fluvic Gleysols (Arenic) and Fluvisol (Arenic) were found (IUSS Working Group WRB, 2022). The plant at Red River is typical of a dry heath tundra, dominated by dwarf shrubs that lose their leaves annually (such as *Betula nana, Vaccinium vitis-idaea, Salix cinerea)* and those that remain green year-round (like *Cassiope tetragona)*. The biocrust layer is composed of lichens, including *Cetraria islandica* and *Stereocaulon paschale* spp. and mosses, such as *Tomentypnum nitens* and *Aulacomnium turgidum* (Xu & Ambus, 2025) but not defined lichens from our sampling site.

The analysis of the soil properties, GHG production rates, and N transformation rates was split between the University of Eastern Finland in Kuopio, Finland, University of Hamburg, and Hereon institute, both in Hamburg, Germany, with some differences in the methods depending on the existing laboratory facilities at each institute. In the supplementary methods, we provide a more detailed description of the methods and an overview table.

#### General soil characteristics

Fresh bulk density was determined by weighing soil samples with a known volume, and dry bulk density (BD) was calculated using the water content. For the determination of water content, soils were dried at 105 °C for 24 hours. Water-filled pore space (WFPS) was calculated according to Haney & Haney. (2010) by WFPS (%) = VWC/(1-BD/PD), where VWC is volumetric water content (%), BD is dry bulk density, and PD is particle density. A constant value of 2.64 g cm-3 was used for our mineral soils. pH and electrical conductivity (EC) were measured from a soil slurry with a soil: $H_2O$ ratio of 1:4. Soil texture, namely %-fractions of sand, silt, and clay, was determined with Sedimat 4-12 (Umwelt-Geräte-Technik GmbH UGT, Müncheberg, Germany). From soils with total C content above 1.5%, humus was removed prior to analysis of soil texture (Van Reeuwijk, 2002).

#### Total, extractable and microbial N and C pools and optical analyses of DOC

The total C and N content was measured with an Elementar VarioMAX Cube Analyzer (Elementar Analysesysteme GmbH, Hanau, Germany). Total inorganic carbon content (TIC) was determined with the Elementar soliTOC Cube Analyzer (Elementar Analysesysteme GmbH, Hanau, Germany) by temperature-dependent differentiation of total carbon as described by Mörchen et al., 2019. From the measured TC and TIC, the total organic carbon (TOC) content was calculated. Total N content was below the detection limit of 0.01% in three samples: RB2 0-1 cm, RB2 1-2 cm, and RB2 13+ cm. For these samples, N content was set to half of the detection limit for data analyses.

For the determination of mineral N content, soils were extracted using a 1:4 soil: extractant ratio (w/v) with either 1 M KCL (Kuopio) or 0.0125 M $CaCl_2$ (Hamburg), depending on the analytical method used for concentration determination. In Hamburg, the nitrite ($NO_2^-$) and $NO_3^-$ concentrations were measured using High-Performance Liquid Chromatography (HPLC) (AA280FS, Agilent, Santa Clara, USA). In Kuopio, $NO_2^-$ and $NO_3^-$ concentrations in the soil extracts were measured colorimetrically utilizing the reaction of $NO_2^-$ with the Griess reagent (Miranda et al., 2001). First, $NO_2^-$ was determined by measuring absorbance at 540 nm using a photometer (DR 5000, Hach Lange, Düsseldorf, Germany) with a Sinergy H1 microplate reader (BioTek® Agilent, California, U.S.). Then, $NO_3^-$ was reduced to $NO_2^-$ for the determination of $NO_2^- + NO_3^-$ by incubating the extracts with VCl3 for 90 min at 32°C. The concentration of $NO_3^-$ was calculated as the difference between the second and the first measurement. In Hamburg, $NH_4^+$ concentration was measured colorimetrically by a method in which indophenol blue is produced by reaction of $NH_4^+$ with salicylate and hypochlorite in the presence of sodium nitroprusside at 655 nm (Urban, 2002). In Kuopio, the method was almost similar with the difference that sodium-phenate was used instead of salicylate and the absorbance was measured at 650 nm (Voigt et al., 2007). We harmonized the results by $CaCl_2$ extractions to KCl extractions by 0.5 according to Hansen & Elberling., 2023).

Dissolved organic carbon (DOC), total dissolved nitrogen (DN), and aromaticity of DOC were measured from water extracts using a 1:4 soil: extractant ratio (w/v). The samples were filtered with 0.45 µl syringe filters before the analysis on the TOC/TN analyzer (Shimadzu TOC-VCPH, Shimadzu Corp., Kyoto, Japan). The aromaticity of DOC, inversely related to its biodegradability, is described by the wavelength-specific UV-absorbance at 254 nm, $SUVA_{254}$, which was measured with a spectrophotometer (UV-1800, Shimadzu, Kyoto, Japan). Calculation of aromaticity followed Weishaar et al. (2003). From the water extracts, we also measured anion concentrations, including chloride ($Cl^-$), sulphate ($SO_4^{2-}$), and phosphate ($PO_3^-$) by ion chromatography (Dionex ICS-2100, Thermo Scientific).

The microbial biomass was determined only from a subset of the Red river samples (RV-2 and RB-2). For this purpose, we used a previously described chloroform fumigation extraction (CFE) method (Brookes et al., 1985, Vance et al., 1987). The samples were incubated for 24 h at 22°C with and without ethanol-free chloroform and extracted with 0.5M K2SO4 with a 1:4 soil: extractant ratio (w/v). The DOC and DN concentrations in filtered extracts were analysed as described above. $C_{mic}$ and $N_{mic}$ were calculated as the difference between chloroform-treated and non-chloroform-treated samples and corrected for incomplete extraction efficiency by dividing the results by published correction factors for C (kEC=0.38) and N (kEN=0.54) (Brookes et al., 1985).

**Soil fluxes**

In situ fluxes from the soil surface were measured using a manual chamber technique using two portable greenhouse gas analyzers (LI-7820 for $N_2O$/$H_2O$ and LI-7810 for $CH_4$/$CO_2$/$H_2O$, LI-COR Biosciences, Lincoln, Nebraska, USA). Transparent, cylindrical plexiglas chambers with a headspace volume of 9.2 L were placed on PVC collars (r = 10 cm), installed more than 24 h in advance. A reflective tarp was used for another measurement under dark conditions. We report fluxes from dark measurements for $N_2O$ (except for plot 2-1, where only light measurements were done) and for $CH_4$. For $CO_2$, we got the net ecosystem exchange (NEE) from the light measurement and ecosystem respiration from the dark measurement, and gross photosynthesis (GPP) was calculated as their difference (GPP=NEE-ER). The chamber was equipped with a fan, a pressure vent, and a thermometer (VOLTCRAFT DET3R, Germany). Gas concentrations were recorded at 1s intervals over a closure time of 3 minutes. Fluxes were calculated based on the concentration change in the chamber headspace over time with linear regression fitting in MATLAB version R2023b using a published chamber flux script (Eckhardt & Kutzbach, 2016), v16 and v17). The first 10 s after chamber closure were discarded. Quality control was based on the root mean square error (RMSE). If the RMSE was higher than 0.4 ppb for $N_2O$, 0.6 ppb for $CH_4$, and 3.5 ppb for $CO_2$, the measurement interval was manually adjusted to ensure stable linear conditions with an RMSE below the set limit. About 5% of the $N_2O$ fluxes were lower than the calculated mean detectable flux but were used to avoid overestimation of $N_2O$ fluxes.

**Water fluxes at the Teno river**

Opaque polyethylene plastic buckets with Styrofoam floats and sampling outlets were used (Gålfalk et al., 2013). Chambers reached approximately 3 cm into the water and their height above water was approximately 13 cm. The area of each chamber was 0.12 $m^2$ and the headspace was 0.009 $m^3$. The closure time for the chamber measurements was 21 min, and the gas sampling was done at 0, 3, 7, 14, and 21 min. Gas samples were taken with 60 ml BD PlastipakTM syringes equipped with a BD Connecta 3-way stopcock valve (Becton, Dickinson and Company, NJ, USA) and injected into pre-evacuated 12-ml Labco Exetainer® vials (Labco Ltd., Lampeter, Ceredigion, UK). Concentrations of $N_2O$, $CH_4$ and $CO_2$ in the gas samples were analyzed using a gas chromatograph (Agilent 7890B, USA) equipped with a GX-271 autosampler (Gilson Inc., USA). On each sampling day, standard gases with known concentrations were measured. The linear regression was fitted to concentrations as a function of time. The regression lines with $r^2 > 0.80$ were accepted. Fluxes were calculated from the concentration increase in the chamber using the equation reported by Erkkilä et al. (2018).

**Greenhouse gas production of $N_2O$, $CO_2$ and $CH_4$ in soil incubations**

The production rates of $N_2O$, $CO_2$ and $CH_4$ in aerobic and anaerobic conditions were determined in incubation experiments at +5°C. For anaerobic incubations, 15 g fresh weight (FW) of homogenized soil with field moisture content was weighed into sterile 120 ml glass flasks, flushed 4-5 times with pure $N_2$ inside a glove bag (Kuopio; $O_2 < 0.05\%$) or in an anaerobic cabinet under anoxic conditions (Hamburg; $O_2 < 0.006\%$) and sealed with 2 cm thick rubber septa and screw caps. For aerobic treatment, laboratory air was used for headspace. Three blank flasks without soil were included for both treatments.

In Kuopio, gas samples were collected for aerobic treatment at six time points (days 0, 1, 2, 3, 6 and 7) and for anaerobic treatment at seven time points (days 0, 1, 2, 3, 6, 7 and 14). We used 60 ml plastic syringes (BD PlastipakTM) equipped with 3-way stopcock valves (BD Connecta; both Becton, Dickinson and Company, NJ, USA) for sampling 25 ml of gas from the headspace which was then injected into pre-evacuated 12 ml glass exetainers (Labco) to create overpressure. Prior to sampling, an equivalent volume of laboratory air (aerobic incubation) or pure $N_2$ (anaerobic incubation) was added to the flask to compensate for the sampled volume. Concentrations of $N_2O$, $CH_4$ and $CO_2$ in the gas samples were analyzed using a gas chromatograph (Agilent 7890B, USA) equipped with a GX-271 autosampler (Gilson Inc., USA).

In Hamburg, both the aerobic and anaerobic flasks were sampled on incubation days 0, 1, 3, 6, 7, 9, 14 and 20. On each sampling day, different standard gases with known concentrations were measured. The standards for $CO_2$ were 101000 ppm, 3990 ppm, 992 ppm, 882 ppm and 347 ppm, for $CH_4$ 100000 ppm, 10000 ppm, 996 ppm, 199 ppm and 1.65 ppm and for $N_2O$ 1.55 ppm, 1.05 ppm and 0.29 ppm. Before each sampling, the pressure within the incubation flasks was measured with a manometer Leo 1 (Keller, Schweiz). In case of under pressure, more N2 (for the anaerobic flasks) or lab air (for the aerobic flasks) was injected into the headspace until overpressure was reached. Samples were taken with a 1 ml syringe with a black needle (Sterican, 0.70 x 30 mm, Braun Medical AG, Melsungen, Germany). The syringe was injected into the septum in the flask and flushed once with the headspace gas. A 1 ml gas sample was taken out of the flask and injected directly into the GC (Agilent 7890 A, Santa Clara, USA), which measured the concentration of the gases $CO_2$, $CH_4$ and $N_2O$. If the $CO_2$ concentration in a flask was higher than 30 000 ppm, the flask was flushed with $N_2$ (for the anaerobic flasks) or laboratory air (for the aerobic flasks). Those flushed samples were re-sampled and re-measured after approximately one hour after flushing (new zero point).

Gas production rates were calculated from concentration changes (ppm) during the incubation using the ideal gas law. For incubations in Kuopio, we accounted for the amount of GHGs added to the flask with the laboratory air in the aerobic treatment as well as for those removed with the samplings in both treatments. In Hamburg, additionally, the pressure in the incubation flask, the water volume in the soil sample, and the solubility of GHG in water were considered in the calculation.

#### $^{15}N$ content in mineral N

Due to the limited soil availability for the Red river site, gross mineralization was not determined for RB-1 and RV-1. Gross nitrification was determined in two batches with slight differences in the methodology: All Teno samples and Red river samples RB-1 and RV-1 were processed in Kuopio, while RB-2 and RV-2 were processed in Hamburg.

### $^{15}N$ in $NH_4^+$

Soil extracts were prepared for $^{15}N$-$NH_4^+$ analysis using the microdiffusion method (Lachouani et al., 2010). Before microdiffusion, the $NH_4^+$ concentrations in the extracts were equalized to a target concentration of 80 μmol N l−1 by spiking (low concentrations) or dilution (high concentrations). For gross mineralization, we used a non-labelled $NH_4^+$ spiking solution with a concentration of 500 μmol N $l^{-1}$ and a 1:5 ratio of sample N: total N. For DNRA, the spiking solution had 200 μmol N $l^{-1,}$ and a 2:5 ratio of sample N: total N was used. Soil extracts with low $NH_4^+$ concentrations (<20 μmol N $l^{-1}$) were excluded from the analysis. For each sample batch (microdiffusion + IRMS run), we included duplicate standards (STDs) with the same concentration as in the samples but with three different at% levels: natural abundance (0.366 at%), 5 at%, and 10 at%. The at% results in the samples measured on the IRMS were then corrected using the STD curve of actual vs. measured at% in the STDs.

**$^{15}$N in $NO_3^-$**

In Kuopio, soil extracts were prepared for $^{15}N$-$NO_3^-$ analyses using the Titanium (III) method (Altabet et al., 2019), where $NO_3^-$ is chemically reduced to $N_2O$ using $TiCl_2$ in the acidic (HCl) media, followed by 15N-$N_2O$ analysis of headspace gas by isotope ratio mass spectrometry (IRMS). The method is, to a certain extent, sensitive to $N_2O$ concentration (Altabet et al., 2019). For the DNRA assay, we equalized the concentrations in the 3 ml of final sample used for the reaction to 14.3 μmol N l-1 by spiking them with non-labelled $NO_3^-$ solution with a concentration of 100 μmol N $l^{-1}$ and a 1:4 ratio of sample N: total N. The final 15N results were then corrected for the dilution caused by spiking. For each sample batch (Ti-method + IRMS run), we included duplicate standards (STDs) with the same concentration as in the samples but with three different at% levels: natural abundance (0.366 at%), 2 at%, and 5 at%. The at% results in the samples measured on the IRMS were then corrected using the STD curve of actual vs. measured at% in the STDs.

In Hamburg, $^{15}$N isotopic composition was measured using bacterial conversion to $N_2O$ *(Ps. aureofaciens),* analyzed by IRMS (Termo) (Sigman et al., 2001; Casciotti et al., 2002). For the calibration, we measured in each run two international standards (USGS-34; IAEA-$NO_3^-$) and one internal standard (Sigman et al., 2001; Casciotti et al., 2002). Cross-comparison of the two analytical methods used for $^{15}N$-$NO_3^-$ determination confirmed very similar results across the two methods and labs.

**Results**

**Figure S1.** Soil horizon profile for the Red River soil samples. RB code represents bare, and RV code represents vegetated samples. Soil horizons according to depth at Red River, with C content (%) shown for each layer.

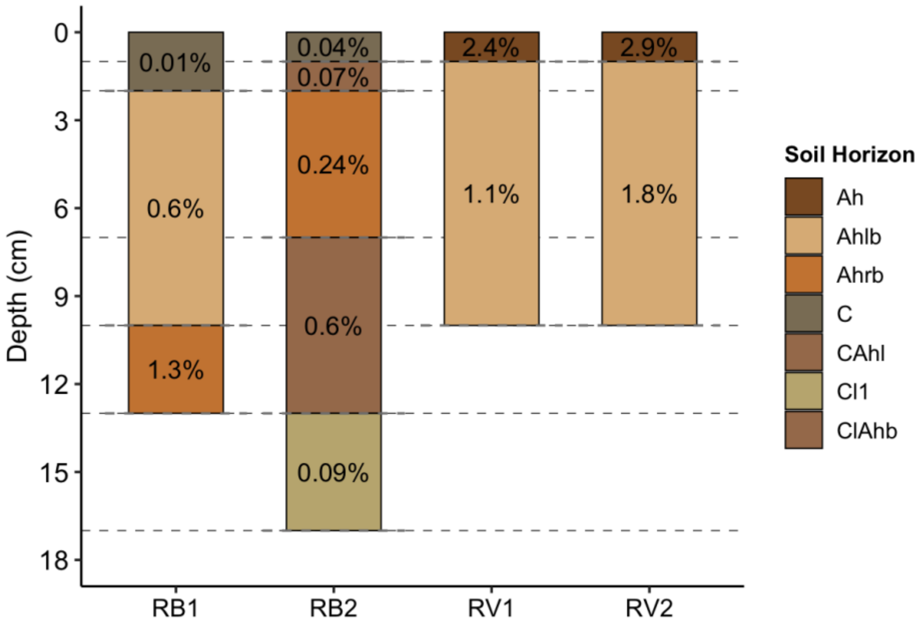

**Figure S2.** Soil profile pictures at the Red River and the Teno River

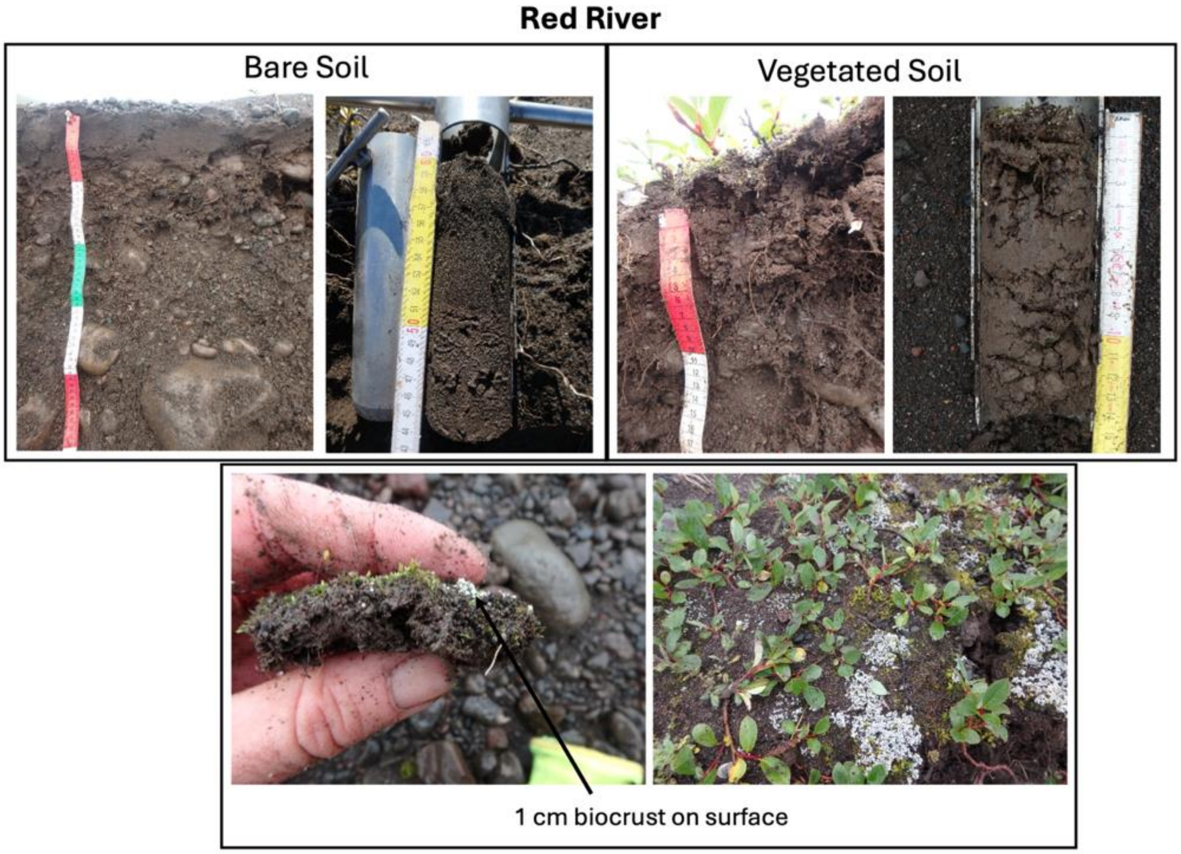

Teno river (shoreline)

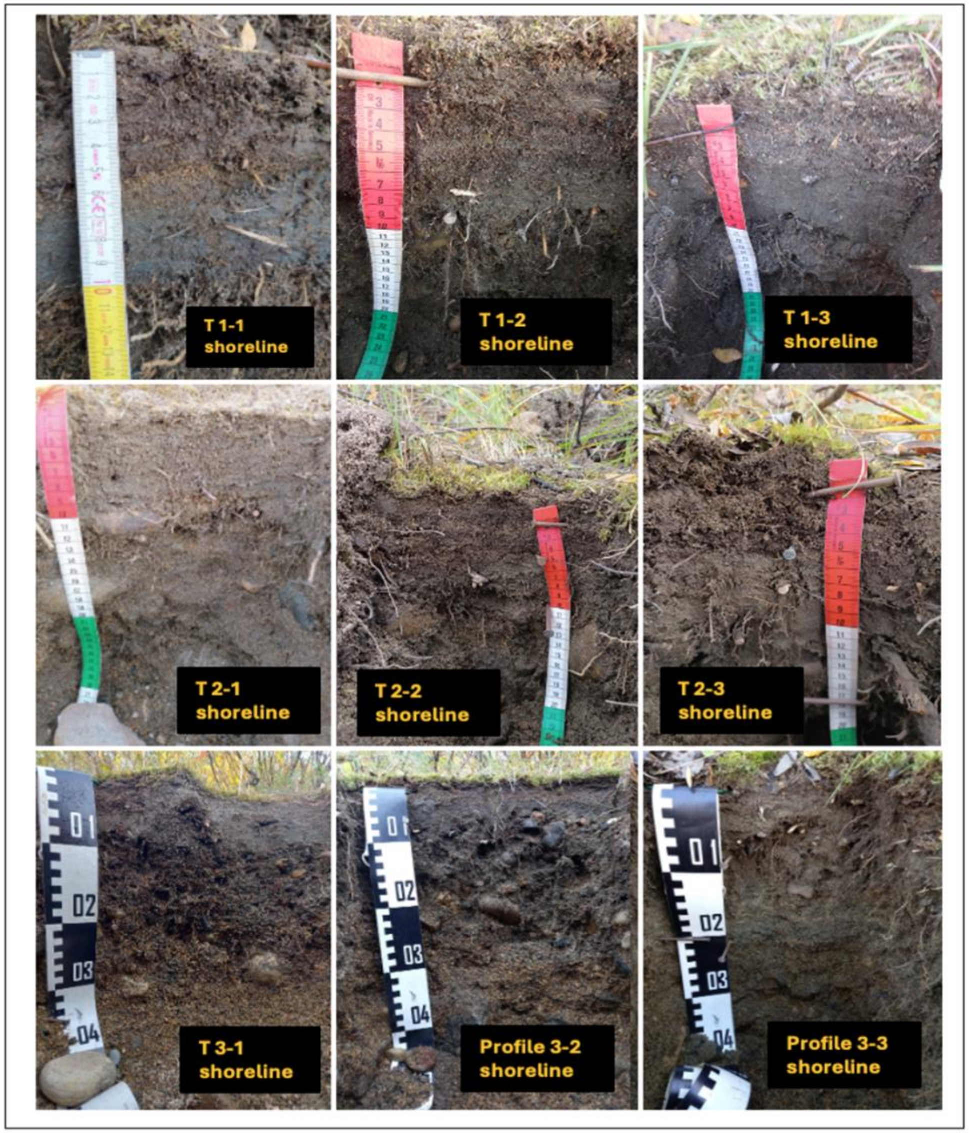


Teno river (Sandbank)

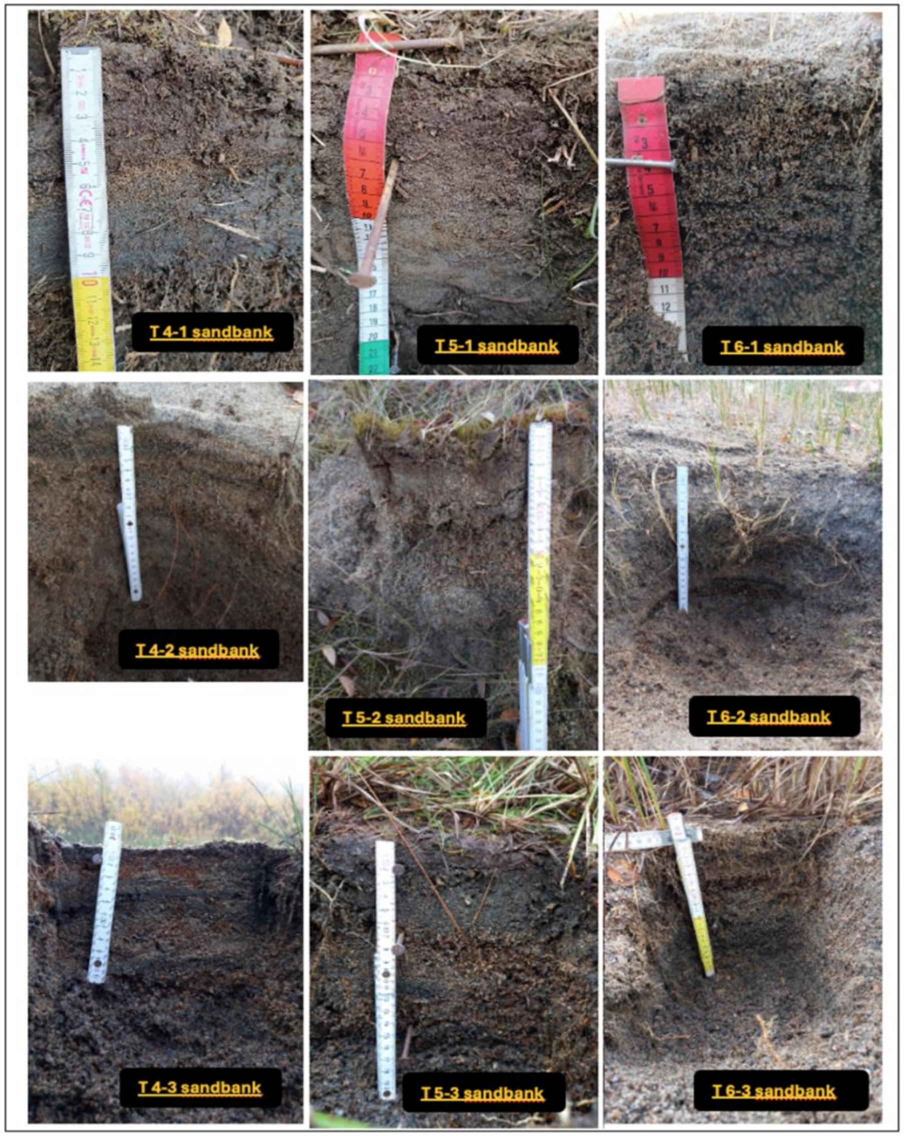

**Figure S3.** Soil characteristics, including WFPS, pH, bulk density, soil texture in bare (brown; B) and vegetated (green; V) soils, ***a)*** at the Teno river along the distance gradient from the shoreline, ***b)*** at Red River along depth profile.

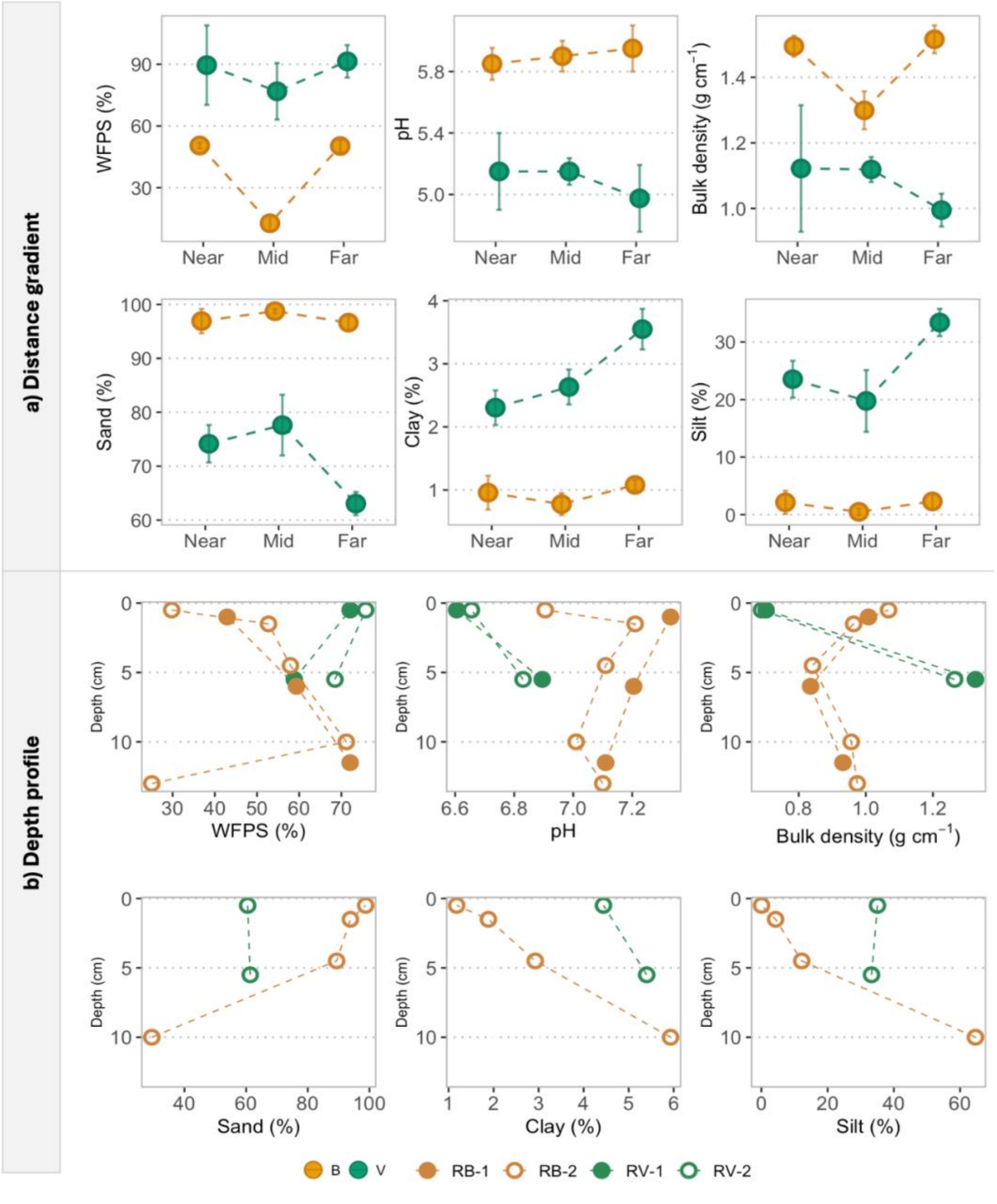

**Figure S4.** Content of various N and C forms, including total C, total N, C/N ratio, DOC, DON and mineral N species in bare (brown; B) and vegetated (green; V) soils at the Red River along depth profiles**.** bare plots are coded as RB and vegetated plots as RV.

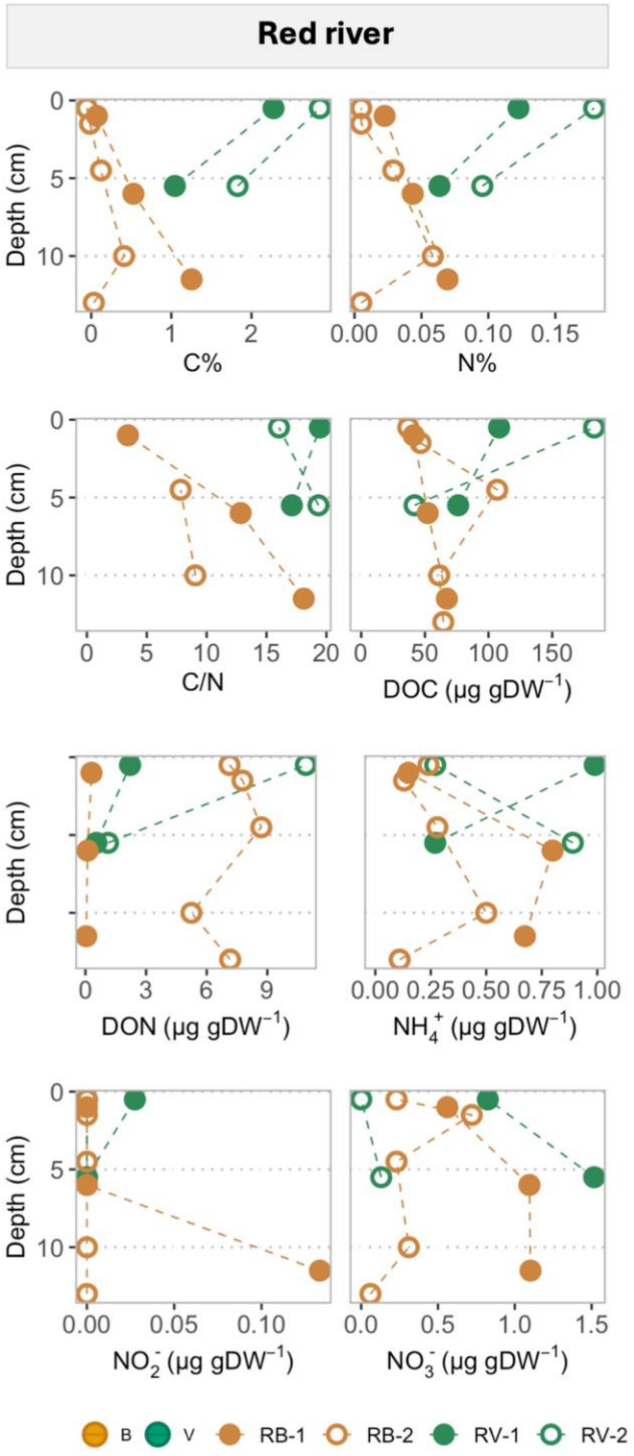

**Figure S5.** Nitrogen transformation rates including net mineralization, net nitrification, gross mineralization, gross nitrification, potential denitrification and potential DNRA, at the Red River in bare (brown; B) and vegetated (green; V) soils, along depth profile.

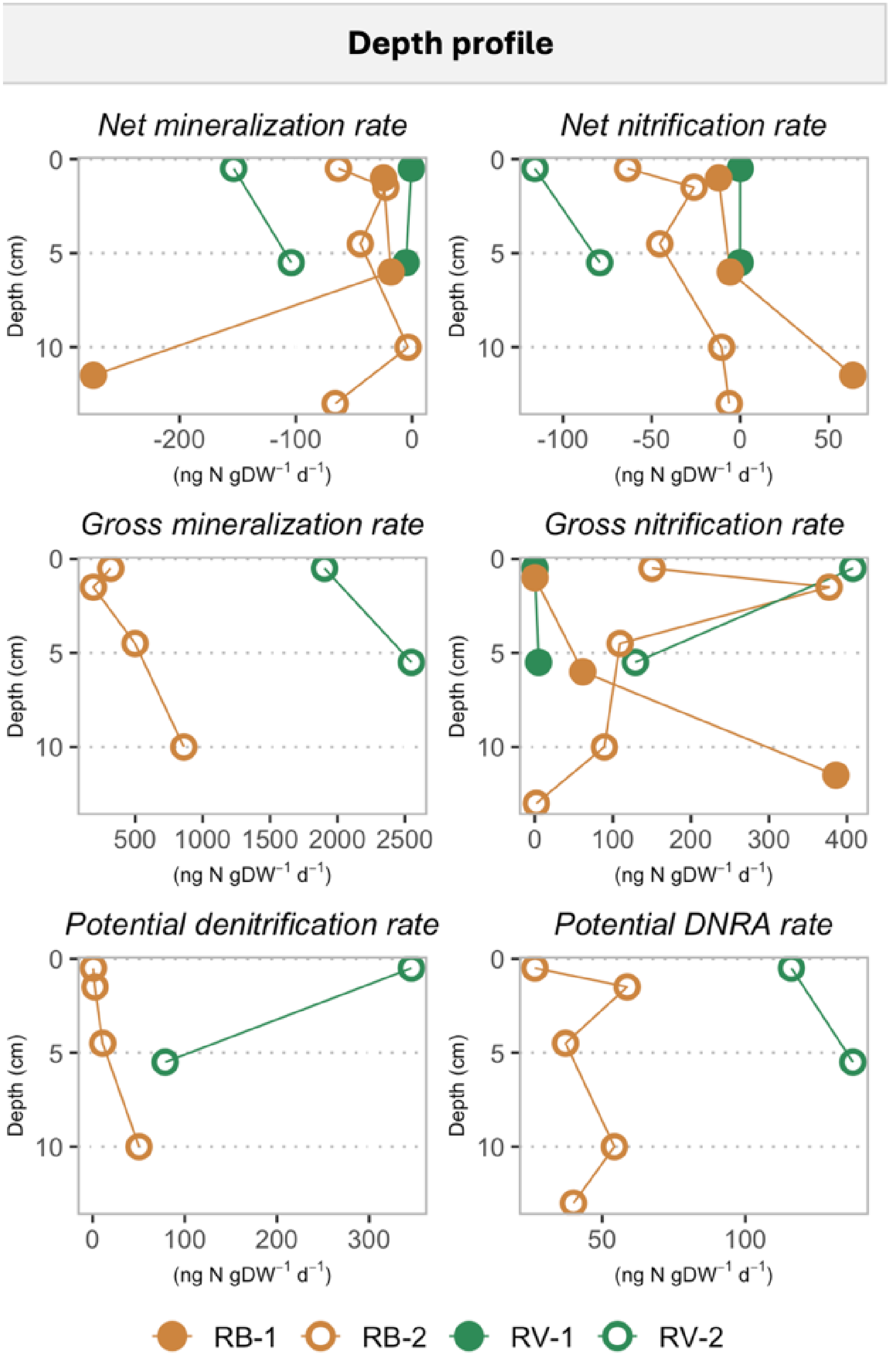

**Figure S6.** ER, NEE and GPP at the Teno River.

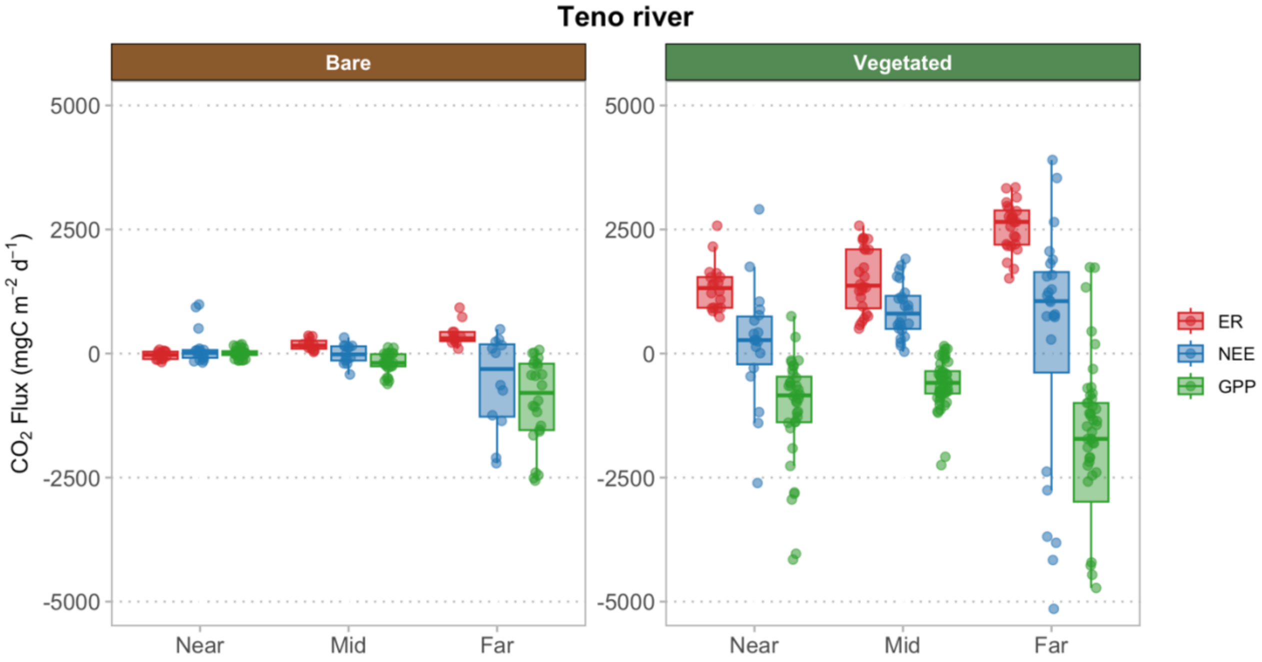


**Figure S7.** Light and dark measurements of $CO_2$ fluxes at the Red River (n=2-4).

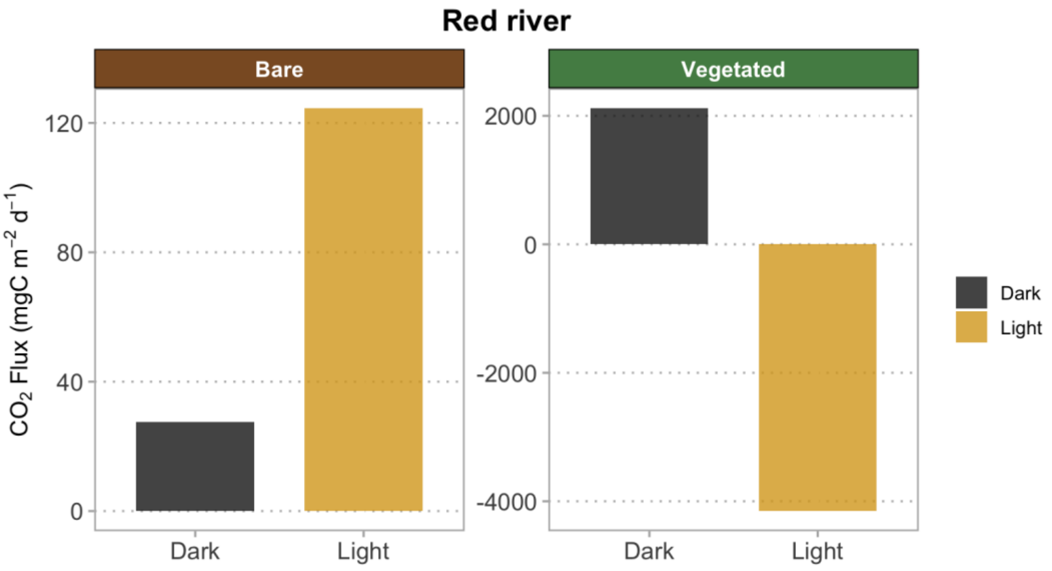

**Figure S8.** Gas production in oxic and anoxic incubations at Red River in bare (brown; B) and vegetated (green; V) soils along the depth gradient. Point plots show mean value.

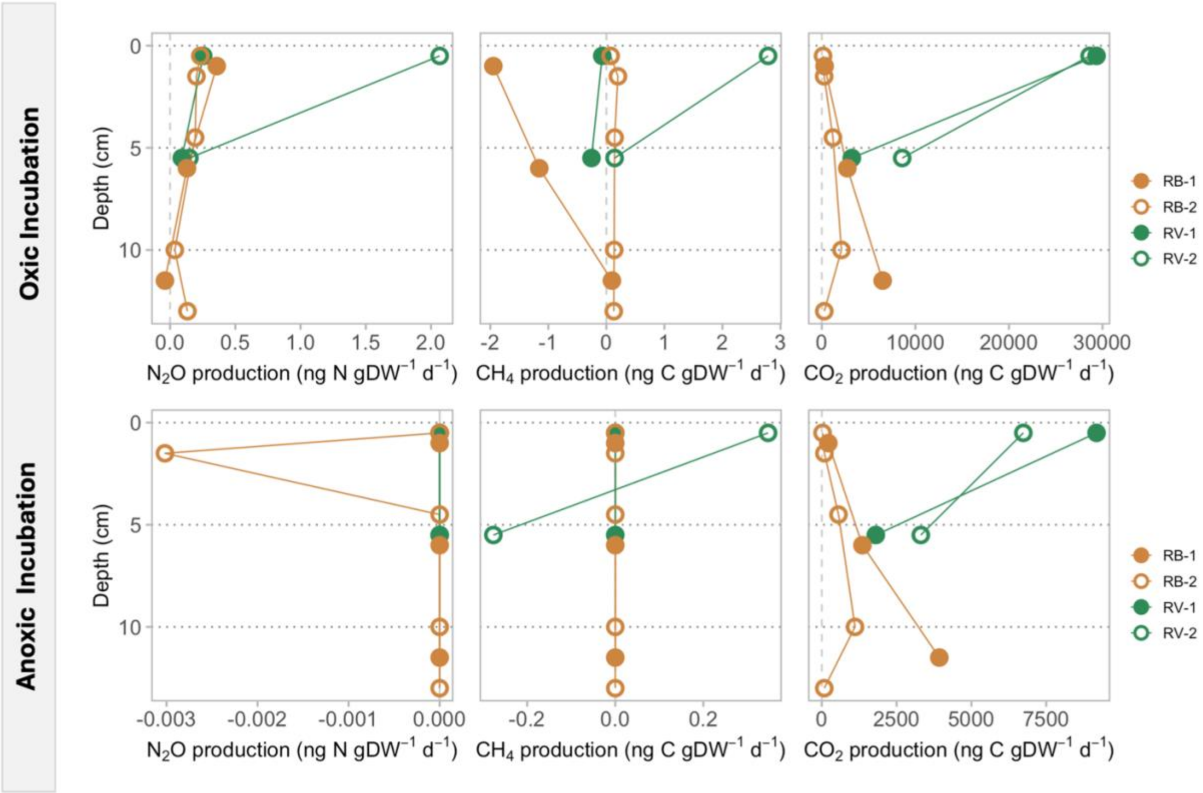

**Figure S9.** Scree plots and contribution plots to PC are shown first four for he Teno river and the bottom four plots for the Red River.

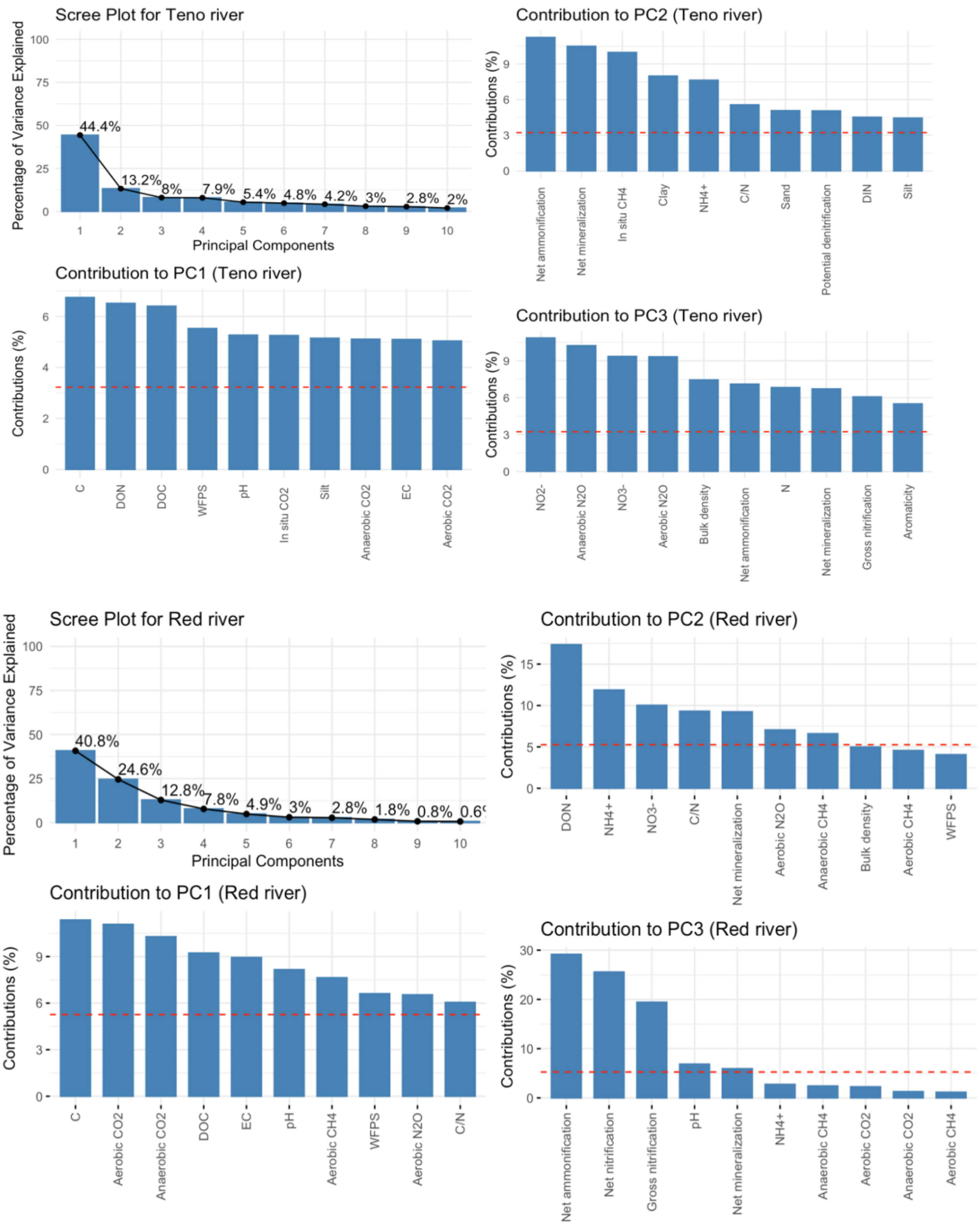

**Figure S10.** Correlation heat map showing the Pearson correlation between N pools, N transformation process, greenhouse gas production, *in situ* field, and soil characteristics at the Teno River. A negative correlation is shown in blue, and a positive correlation is shown in red.

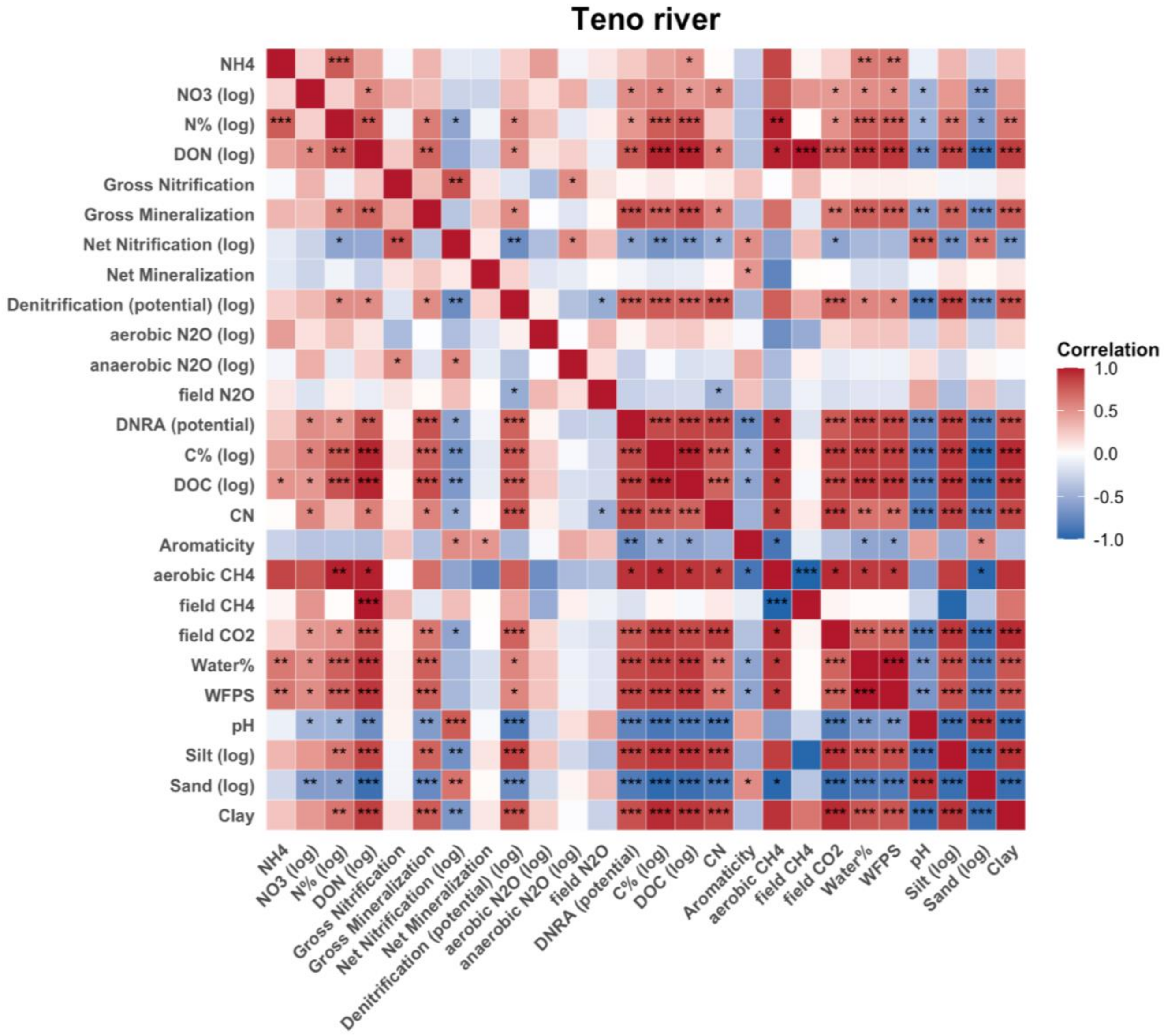

**Figure S11.** Correlation heat map showing the Pearson correlation between N pools, N transformation process, greenhouse gas production, and soil characteristics at the Red River. A negative correlation is shown in blue, and a positive correlation is shown in red.

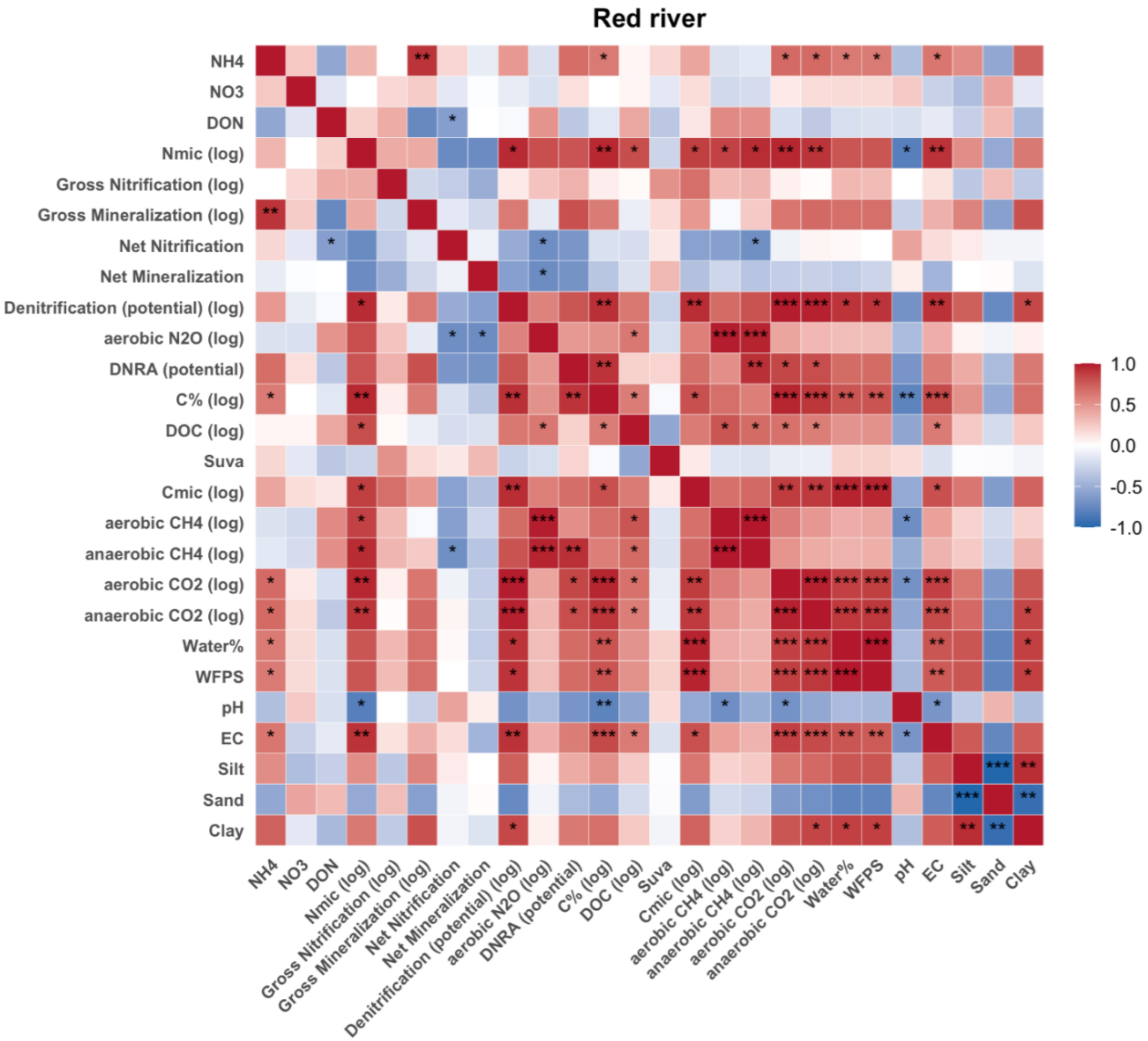

**Figure S12.** Gross mineralization rates measured in this study in comparison to those predicted using published regression functions from a meta-analysis on permafrost-affected soils (Ramm et al. 2022). a) Comparison of rates predicted based on SOC content; b) Comparison of rates predicted based on N content; c) overall magnitude of mineralization with the predicted SOC and N content.

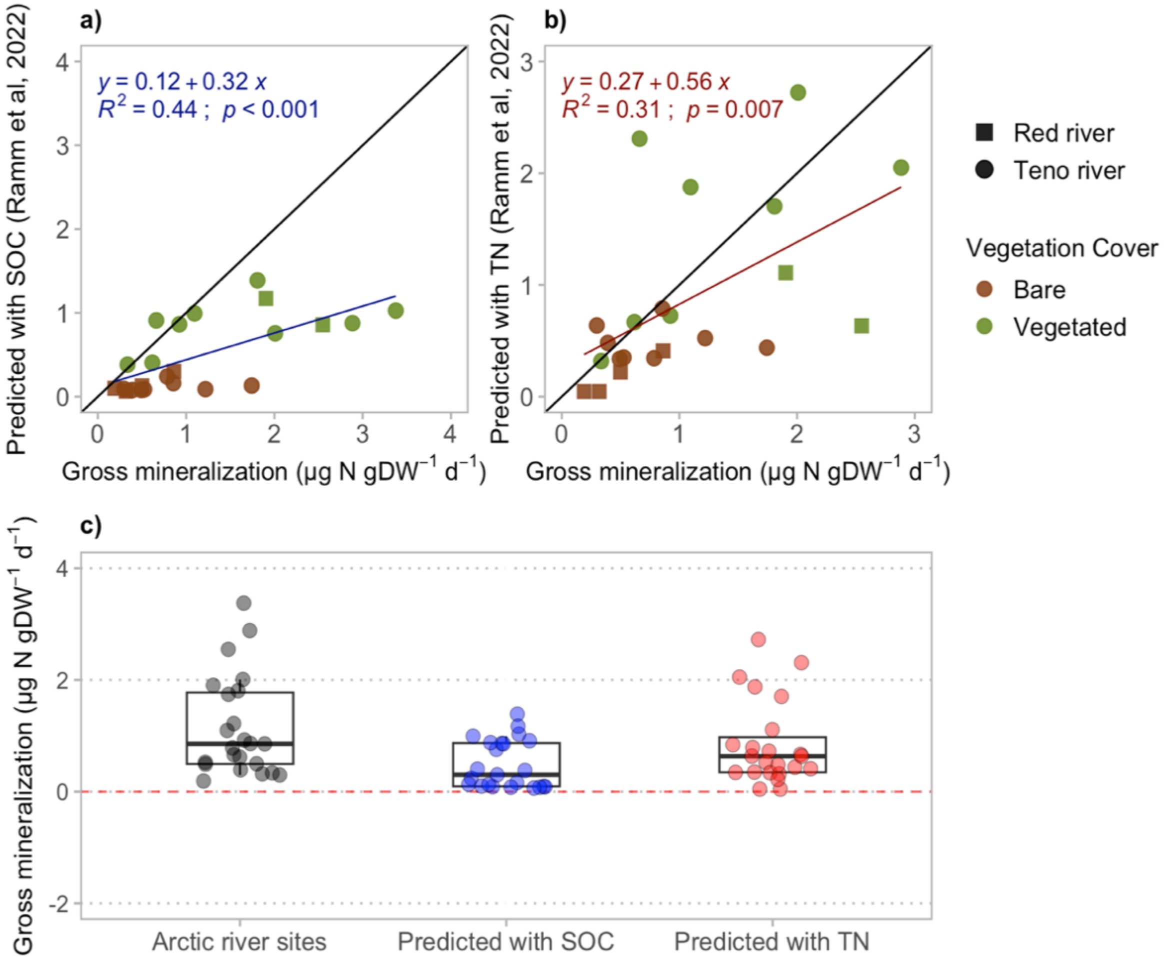

| | Teno River | | | Red River | | |
|---|---|---|---|---|---|---|
| | Bare (n= 8) | Vegetated (n= 10) | | Bare (n= 24) | Vegetated (n= 12) | |
| Sand (%) | 97.3 ± 3.2 | 72.0 ± 10.1 | *** | 81.0 ± 29.1 | 60.9 ± 0.6 | ns |
| Silt (%) | 1.8 ± 2.9 | 25.1 ± 9.6 | *** | 16.3 ± 24.3 | 34.2 ± 1.3 | ns |
| Clay (%) | 2.9 ± 0.8 | 3.7 ± 2.2 | *** | 1.2 ± 0.7 | 3.1 ± 1.7 | ns |
| Bulk Density (g $cm^{-3}$) | 1.5 ± 0.1 | 1.1 ± 0.1 | *** | 0.9 ± 0.1 | 1.0 ± 0.3 | ns |
| C (%) | 0.1 ± 0.1 | 2.0 ± 0.9 | *** | 0.4 ± 0.4 | 2.2 ± 0.8 | *** |
| N (%) | 0.07 ± 0.03 | 0.3 ± 0.2 | ** | 0.04 ± 0.02 | 0.1 ± 0.05 | *** |
| C/N ratio | 2.1 ± 1.9 | 9.6 ± 5.3 | ** | 11.1 ± 5.1 | 18.7 ± 1.6 | *** |
| DOC (µg $gDW^{-1}$) | 49 ± 13 | 313 ± 145 | *** | 60 ± 30 | 102 ± 59 | *** |
| DOC aromaticity (%) | 1.0 ± 0.01 | 1.1 ± 0.8 | ns | 1.1 ± 0.04 | 1.7 ± 0.36 | ns |
| DON (µgN $gDW^{-1}$) | 0.2 ± 0.4 | 7.0 ± 4.1 | *** | 2.2 ± 3.0 | 4.5 ± 5.1 | ns |
| $NH_4^+$ (µgN $gDW^{-1}$) | 1.5 ± 1.2 | 3.4 ± 2.1 | * | 0.4 ± 0.2 | 0.6 ± 0.3 | ns |
| $NO_2^-$ (µgN $gDW^{-1}$) | 0.02 ± 0.03 | 0.1 ± 0.1 | * | bd | bd | - |
| $NO_3^-$ (µgN $gDW^{-1}$) | 0.3 ± 0.1 | 1.2 ± 1.0 | * | 0.1 ± 0.05 | 0.1 ± 0.07 | ns |
| Cmic (µg $gDW^{-1}$) | na | na | - | 539.0 ± 352 | 1949.1 ± 1174 | ** |
| Nmic (µg $gDW^{-1}$) | na | na | - | 6.2 ± 7.3 | 135.1 ± 128.5 | ns |
| WFPS (%) | 41.0 ± 17.8 | 85.2 ± 21.7 | *** | 54.3 ± 15.6 | 67.1 ± 7.1 | * |
| Water (%) | 15.5 ± 6.7 | 32.2 ± 8.2 | *** | 19.5 ± 7.0 | 25.7 ± 3.0 | ** |
| pH | 5.9 ± 0.2 | 5.1 ± 0.3 | *** | 7.1 ± 0.1 | 6.7 ± 0.1 | *** |
| EC (µS $cm^{-1}$) | 13.4 ± 8.7 | 27.6 ± 9.8 | ** | 19.6 ± 7.8 | 28.9 ± 6.7 | ** |
| $Cl^-$ (µg $gDW^{-1}$) | 4.6 ± 9.4 | 9.8 ± 5.8 | ns | 0.7 ± 0.5 | 2.8 ± 2.6 | * |
| $SO_4^{2-}$ (µg $gDW^{-1}$) | 1.7 ± 1.4 | 5.7 ± 10.0 | ns | 0.5 ± 0.5 | 2.1 ± 1.0 | ** |
| $PO_4^{3-}$ (µg $gDW^{-1}$) | bd | bd | - | bd | bd | - |

**Table S1.** Soil characteristics in the riparian zones of Teno and Red rivers. Physical and chemical properties are shown for bare and vegetated soils, including soil texture, bulk density, N%, C%, C/N ratio, water filled pore space (WFPS), water content, pH, electrical conductivity (EC), microbial biomass C ($C_{mic}$) and N ($N_{mic}$), and extractable chloride and sulphate content. Data is presented as means with standard deviations. T-test significance is shown by *** = $p < 0.001$, ** = $p < 0.01$, * = $p < 0.05$, and ns = not significant. b.d. = below detection limit.
